# *De novo* design of mini-proteins as specific binders to PACAP-38, a key mediator of migraine

**DOI:** 10.64898/2026.10.30.763049

**Authors:** Hiroki Yamaguchi, Shota Suzuki, Yoshinori Fujiyoshi, Mark Isalan

**Affiliations:** Department of Life Sciences, Imperial College London, Exhibition Road, SW7 2AZ London, UK; Research Institute for Bioscience Products & Fine Chemicals, Ajinomoto Co., Inc. 1-1 Suzuki-cho, Kawasaki-ku, Kawasaki 210-8681, Japan; Advanced Research Initiative, Institute of Integrated Research, Institute of Science Tokyo, 1-5-45 Yushima, Bunkyo-ku, Tokyo, 113-8501, Japan; CeSPIA Inc., 2-1-1 Otemachi, Chiyoda-ku, Tokyo, 100-0004, Japan; Joint Research Course for Advanced Biomolecular Characterization, Faculty of Agriculture, Tokyo University of Agriculture and Technology, 3-5-8 Saiwai-cho, Fuchu, Tokyo, 183-8509, Japan; Imperial College Centre for Engineering Biology, Imperial College London, London, SW7 2AZ, UK

**Keywords:** *De novo* design, PACAP-38, Bindcraft, ProteinMPNN, Migraine, Protein engineering

## Abstract

Pituitary adenylate cyclase-activating peptide 38 (PACAP-38) is a vasoactive peptide that is attracting attention as a potential target for next-generation migraine treatments. Antibodies against PACAP-38 have recently shown promising results as a ligand trap medicine in clinical trials. In this study, we investigated the *de novo* design of miniproteins to evaluate the potential for developing a medium-molecular-weight drug that binds to PACAP-38. Using Bindcraft, we designed binders against PACAP-38 and identified ICLIST-2, a 63-residue peptide with affinity for PACAP-38 (*K*_D_ = 26.5 nM). Furthermore, by alanine scanning and structural analysis of ICLIST-2, we determined residues that were suitable for further optimization using ProteinMPNN. This resulted in the design of ICLIST-11, which incorporates nine amino acid substitutions relative to ICLIST-2, resulting in a *K*_D_ of 4.51 nM. Interaction analysis based on a structural model suggested that three mutations in ICLIST-11 may be important for enhancing affinity. Moreover, ICLIST-11 exhibits a *T*_m_ value that is 14 °C higher, and MD simulations suggest that the substitution of three residues with hydrophobic residues (E30F, M31L, and R54W) is one reason for this. In cell-based functional assays, ICLIST-11 demonstrated potent, concentration-dependent neutralizing activity against PACAP-38-induced PAC1R activation (IC50 = 10.6 nM), significantly outperforming the parental ICLIST-2. Taken together, these results provide a framework for protein drug discovery targeting PACAP-38.

## Introduction

Migraine is a chronic neurological disorder that is estimated to affect around one billion people worldwide^1^. It is the second most common neurological disorder after tension-type headache, with a reported female-to-male sufferer ratio of 3:1 and a prevalence peak between the ages of 35 and 39^1, 2^. According to the clinical criteria, migraine is defined as recurrent headaches lasting 4–72 hours, accompanied by moderate to severe pain^3^. Consequently, migraine significantly impairs patients’ quality of life and results in considerable social and economic costs. While the exact mechanisms triggering migraine attacks remain unclear, the early stages of onset are believed to involve the activation and sensitization of first-order trigeminovascular neurons, as first proposed by Moskowitz in 1984^4, 5^. This leads to the release of vasoactive peptides, such as calcitonin gene-related peptide (CGRP) and pituitary adenylate cyclase-activating peptide 38 (PACAP-38), as well as nitric oxide. This induces a local inflammatory response^6^.

Over the past few decades, the greatest paradigm shift in migraine drug development has been the emergence of CGRP-targeting therapies^7^. CGRP is a signaling molecule with potent vasodilatory effects, and the development of small-molecule CGRP receptor antagonists, as well as monoclonal antibody drugs that target CGRP and its receptors, has transformed the prevention and treatment of migraine^7^. However, a major clinical challenge remains: even with these new CGRP inhibitors, approximately 40–50% of patients still exhibit ‘treatment resistance’, failing to achieve an adequate therapeutic response^8, 9, 10, 11^. This strongly suggests the existence of an additional signaling pathway in migraine pathophysiology that is independent of, yet complementary to, the CGRP pathway^12^. Against this backdrop, PACAP-38 is currently attracting the most attention as a next-generation therapeutic target. PACAP-38 is a 38-amino acid neuropeptide isolated from ovine hypothalamic tissues in 1989^13^; like CGRP, it has potent vasodilatory effects and intravenous administration has been shown to reliably induce migraine-like attacks in patients^7^. Furthermore, recent clinical studies have revealed that these PACAP-38-induced attacks are not suppressed by blocking the CGRP pathway with CGRP antibodies prior to administration, thereby proving that PACAP-38 drives a pathophysiological pathway that is completely independent of CGRP^14^. PACAP-38 activates three class B G protein-coupled receptors (GPCRs): the pituitary adenylate cyclase-activating polypeptide type I receptor (PAC1R); the vasoactive intestinal polypeptide receptor 1 (VPAC1R); and the vasoactive intestinal polypeptide receptor 2 (VPAC2R)^15^. PAC1R is a PACAP-selective receptor, while VPAC1R and VPAC2R have similar affinities for PACAP-38 and vasoactive intestinal peptide (VIP)^13, 16^. Based on the hypothesis that PACAP triggers migraine by activating PAC1R^17^, monoclonal antibodies targeting PAC1R have been developed in recent years. However, they failed to demonstrate a clear effect at the proof-of-concept stage^18^. This suggests that the clinical mechanism of PACAP has not yet been fully elucidated. Nevertheless, bocunebart (Lu AG09222), a monoclonal antibody that specifically inhibits PACAP-38, has been shown to significantly reduce the number of migraine days per month, while demonstrating an excellent safety profile, in clinical trials involving patients with treatment-resistant migraine^19, 20^. The success of bocunebart clearly demonstrates the clinical efficacy of targeting PACAP-38.

However, as antibody therapeutics become more widespread, their limitations are also becoming apparent. As biological agents, antibodies are extremely costly to manufacture, and the associated healthcare costs and financial burden on patients are major practical barriers to their widespread use. Furthermore, structural challenges remain, such as the restriction to injection as the route of administration, and reduced tissue permeability due to their large molecular weight. One approach to overcoming these limitations is peptide therapeutics, which combine the advantages of antibodies and small molecules^21, 22^. As the interaction between PACAP-38 and PAC1R forms a relatively broad protein-protein interaction interface^23^, inhibition using conventional small-molecule compounds was thought to be difficult. However, appropriately designed peptide binders can ensure an affinity and specificity equivalent to that of antibodies. A key feature of this modality is that it is a more stable pharmaceutical that can be produced via chemical synthesis or microbial expression. This dramatically reduces transportation, storage and manufacturing costs compared to antibodies. This therefore has the potential to become a therapeutic strategy that offers socio-economic advantages and the possibility of developing simple, less invasive administration formats^21, 22^. In this study, our aim was to create next-generation seed molecules for migraine treatment that block the PACAP-38 signal by attempting the rational design of novel peptide binders with high affinity and selectivity for PACAP-38. This work reports on the screening of peptide binders designed using a computational approach based on structural information, as well as the evaluation of their binding affinities and inhibitory activities against PACAP-38.

## Results and discussion

We aimed to develop PACAP-38 binding peptides, which have ligand trapping function for migraine treatment, based on structural information. The three-dimensional structure of the PACAP-38–antibody complex has not yet been reported. Therefore, we began our design work based on the cryo-EM structure of PAC1/PACAP-38 (PDB ID: 6M1I), as described in previous literature^23^. In this structure, the 11 C-terminal residues (residues 28– 38) of PACAP-38 were highly flexible and disordered, and no model had been constructed for this region. *In vivo*, PACAP exists as both PACAP-38, which comprises 38 residues, and PACAP-27, which is a cleaved form consisting of the N-terminal 27 residues^16^. The amino acid sequence of the first 27 residues of PACAP-38 is highly conserved across almost all vertebrate species and exhibits physiological activity as a functional peptide^24, 25, 26, 27^. As PACAP-27 shares 68% sequence identity with VIP (which consists of 28 residues), PACAP belongs to the glucagon/gastric inhibitory polypeptide (GIP)/secretin/VIP family. A member of this family is involved in regulating the class B G protein-coupled receptor (GPCR) secretin receptor^28^. When designing the binder, we defined the initial target as the N-terminal 27 residues for two reasons. Namely: (i) these 27 residues constitute the minimal unit responsible for sufficient biological activity, and targeting them is expected to effectively block PACAP-38 binding to its receptor; and (ii) the highly flexible C-terminal 11 residues may complicate the design of a specific binder.

Based on these considerations, our top priority was to identify a binder that interacts strongly with the N-terminal 27 residues that are recognised most precisely by the PAC1 receptor. Using Bindcraft^29^, we designed a total of 100 binder candidates, ranging from 31 to 100 residues, based only on the three-dimensional structure of these 27 residues (Fig. 1).

**Fig. 1.**
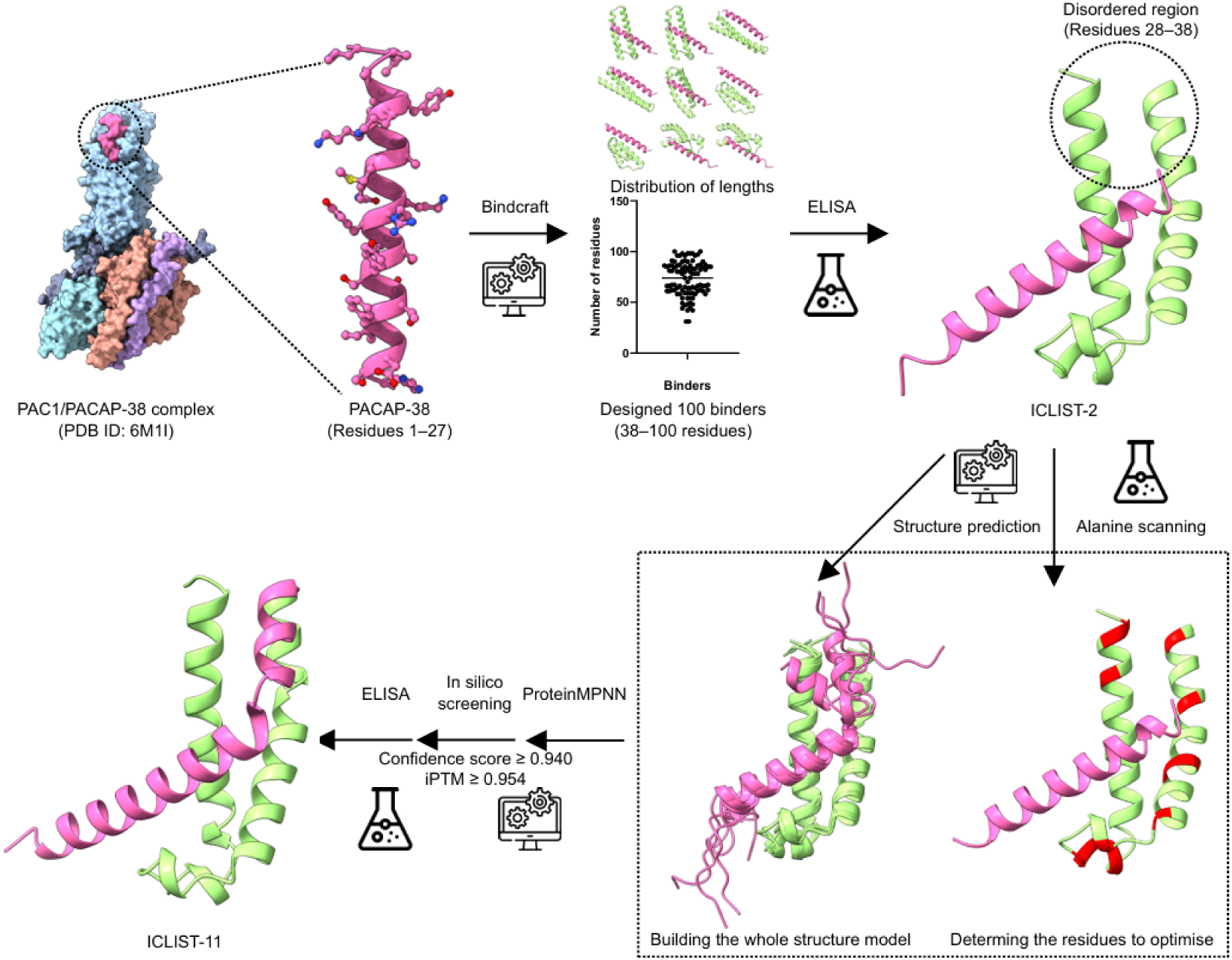
Workflow of PACAP-38 binder design and screening. This workflow outlines the design of PACAP-38 binders in this study. The three-dimensional structure of PACAP-38 (magenta) was extracted from the cryo-EM structure of the PAC1/PACAP-38 complex (PDB ID: 6M1I)^23^. In this PACAP-38 structure, the C-terminal residues (residues 28–38) are disordered. The three-dimensional structures of the binders designed using Bindcraft are shown in green, along with the length distribution of the 100 binders (top centre). The candidate mutant residues identified by alanine scanning of ICLIST-2, which was discovered through ELISA screening, are shown in red (bottom right).

### *De novo* design using Bindcraft

Considering the ease with which the final peptide could potentially be synthesised, our aim was to design a shorter binder. However, we were unable to identify the conditions under which the calculations would converge for peptide designs of 30 residues or fewer. However, on the Ariax bio platform, which was used for the Bindcraft design, we were able to design potential viable binders by setting the parameters to ‘4-stage multimer (hard target)’.

When the top 20 Bindcraft-scored binders (ranging in length from 61 to 96 residues) were evaluated for binding to PACAP-38 by ELISA, only three showed binding (Fig. 2A). These were designated ICLIST-1, -2 and -3, and had lengths of 88, 63 and 63 residues, respectively (Table S1). Furthermore, examining the length distribution of the 100 designed binders revealed 44 proteins were 80 residues or fewer in length, which would be desirable for therapeutic development (Fig. 1). To verify the presence of smaller binders of PACAP-38 not included in the top 20, the binding of these 44 predicted binders, ranging from 31 to 80 residues (excluding the top 20), was assessed by ELISA. However, no binders showing a significant increase in response compared to the control were observed (see Fig. S1). Therefore, the hit rate for the 64 binders evaluated experimentally in this study, excluding duplicates, was estimated to be approximately 4.7%.

**Fig. 2.**
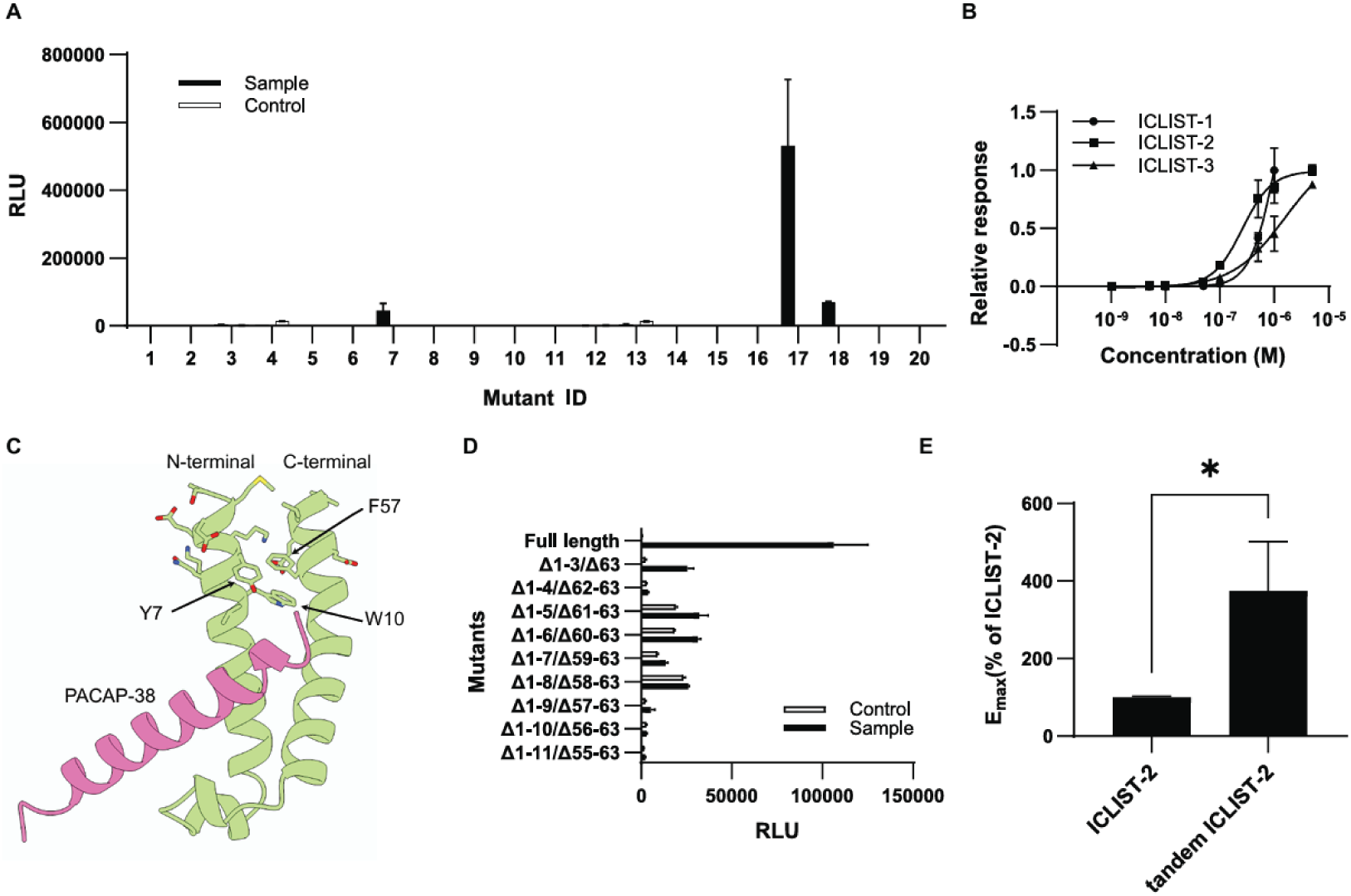
Screening of binders designed by Bindcraft. **A** Binding responses of the top 20 Bindcraft-designed binders for PACAP-38 in ELISA. The control shows a sample without PACAP-38.**B** Concentration-dependent ELISA binding responses of ICLIST-1 (ID 7), ICLIST-2 (ID 17) and ICLIST-3 (ID 18), for a fixed concentration of PACAP-38 (0.1 μg/mL). Fitting was performed using GraphPad Prism 10.6.1. The figures show the relative response, with the binding response at the highest concentration of each binder set to 1. **C** The predicted structure of the complex between ICLIST-2 (green) and PACAP-38 (residue numbers 1–27, pink). **D** Binding responses of N-terminal and C-terminal truncations of ICLIST-2 binding to PACAP-38. The control represents wells without added PACAP-38. **E** E_max_ of tandem ICLIST-2, with ICLIST-2 set to 100%. The experiment was performed three times and the mean ± S.D. is shown.

The EC_50_ values of ICLIST-1, ICLIST-2 and ICLIST-3 for PACAP-38, as measured by ELISA, were 1.0 μM, 258.8 nM and 1.6 μM, respectively (Fig. 2B). ICLIST-2 exhibited the strongest apparent binding response. Furthermore, when affinity for PACAP-38 was assessed using surface plasmon resonance (SPR), ICLIST-2 exhibited the strongest binding with a *K*_D_ of 26.5 nM (see Table 1 and Fig. S2). In this study, a discrepancy of approximately a factor of 10 was observed between the K_D_ values obtained by SPR and the EC_50_ values obtained by ELISA. It is widely recognised that, due to differences in immobilisation methods and measurement principles, the values obtained by ELISA and SPR do not always correspond exactly. The main factors contributing to this discrepancy include the dissociation of the binder during the washing step in ELISA, and steric hindrance associated with physical adsorption onto the plate. Consequently, it is considered that the affinity of this binder, as determined by SPR, is 25 nM, and the ELISA data are valid for screening and relative comparisons. Both ICLIST-2 and ICLIST-3 consist of 63 residues, with seven residues differing between the two (see Fig. S3). Interestingly, six of these residues face outwards rather than towards the PACAP-38 binding surface in the predicted structure, exposing them to the solvent. This suggests that they do not interact directly with PACAP-38, but are likely involved in the binder’s physical properties. Conversely, the side chain of Q8 is the only residue positioned towards the C-terminus of PACAP-38 (residue numbers 1–27). As the disordered region may come into contact with it, Q8 is likely to be a key residue in binding to PACAP-38.

**Table 1.** Parameter of ICLISTs by SPR.

|  | ICLIST-1 | ICLIST-2 | ICLIST-3 | ICLIST-11 |
|---|---|---|---|---|
| *k*<sub>a</sub> (M<sup>-1</sup>s<sup>-1</sup>) | n.d. | 4.1×10<sup>5</sup> | 9.4×10<sup>4</sup> | 9.8×10<sup>5</sup> |
| *k*<sub>d</sub> (s<sup>-1</sup>) | n.d. | 1.1×10<sup>-2</sup> | 1.3×10<sup>-2</sup> | 4.4×10<sup>-3</sup> |
| *K*<sub>D</sub> (nM) | 1000 | 26.5 | 140.1 | 4.5 |

The χ<sup>2</sup> values for the fit were 0.016, 0.226, 1.78 and 1.53 RU, respectively. As ICLIST-1 exhibited a box-shaped sensorgram, making kinetic analysis difficult, an affinity analysis was performed; consequently, no values for *k<sub>a</sub>* or *k<sub>d</sub>* were obtained (indicated as n.d.).

Further investigations were next conducted to verify whether smaller binders could be created based on ICLIST-2. Deletion mutants were evaluated for this purpose. As the region in the vicinity of PACAP-38 was deemed important for α-helix recognition, nine types of truncated variant were prepared with N- and C-terminal deletions. ICLIST-2 was found to have α-helices at both the N- and C-termini, which form a two-helix bundle. Aromatic side chains, such as Y7, W10 and F57, are located at the interface of this bundle (Fig. 2C). Based on this predicted structure, variants were designed in which both the N- and C-terminal regions were truncated at nearly equal intervals. ELISA results showed that truncating just three residues from the N-terminus and one residue from the C-terminus reduced the ELISA response by 76 per cent, and that the response did not significantly recover with shorter variants (see Fig. 2D). Based on these results, we concluded that preparing a truncated form of ICLIST-2, while maintaining its binding to PACAP-38, would be difficult. This is because the α-helices at the N- and C-termini interact tightly via multiple hydrophobic residues, so even a slight truncation could potentially disrupt ICLIST-2 folding. Subsequent studies therefore used a 63-residue ICLIST-2 as a template. Since a binder for the N-terminal 27 residues of PACAP was obtained, a similar design approach could potentially be applied to peptides in the GIP/secretin/VIP family, including VIP.

When designing molecular binders, utilizing the avidity effect through multimerization is an established strategy for enhancing binding affinity and selectivity. Inspired by this approach, we designed and synthesized a new ICLIST “dimer” targeting PACAP-38 to enhance the functionality of ICLIST-2, and verified whether a similar avidity-driven improvement could be achieved within our system. When two copies of ICLIST-2 were linked via a GS linker ((GGGGS)_4_), the resulting construct exhibited good expression and could be purified in the same way as ICLIST-2. The EC50 in the ELISA was 258.8 nM for ICLIST-2 and 141.6 nM for the tandem ICLIST-2, representing a roughly two-fold increase in affinity. The E_max_, with ICLIST-2 set at 100%, was 375% (see Fig. 2B and 2E). Furthermore, when the binding of this tandem ICLIST-2 to PACAP-38 was evaluated by SPR under the same conditions as for single ICLIST-2, the *K*_D_ obtained using the bivalent analyte model was 7.32 nM (Table S2). As a reminder, single ICLIST-2 had weaker binding, with *K*_D_ 26.5 nM (see Table 1). Dissociation was also clearly slower than for single ICLIST-2, suggesting that binding affinity was enhanced by the avidity effect (see Fig. S2B and S2D). These findings suggest that ICLIST-2, like typical binders, is functional as a covalent dimer, and that its binding affinity can be enhanced through the avidity effect.

### Structural validation and alanine scanning

To evaluate the accuracy of the predicted ICLIST-2 structure designed using Bindcraft, ICLIST-2 was overexpressed in *E. coli* and purified. Circular dichroism (CD) measurements were then performed, yielding a spectrum typical of an α-helix. It was estimated that 93.5% of the structure consisted of α-helices (Fig. S4A). This is in very good agreement with the fact that the predicted structure of ICLIST-2 consists almost entirely of α-helices (Fig. 2C). Based on the ICLIST-2/PACAP-38 complex structure, nine residues located within 2.6 Å of the ICLIST-2–PACAP-38 interface were selected and the ΔΔG upon substitution with alanine was calculated using FoldX software^30, 31^ (Fig. 3A). All residues yielded positive ΔΔG values, suggesting that they are important for interacting with PACAP-38. Indeed, when evaluating binding of these alanine replacement mutants to PACAP-38, binding was significantly reduced in eight mutants, with the exception of I46A. I46A involves a substitution from the amino acid isoleucine to alanine and is presumed to have had little effect on PACAP-38 recognition because both residues are neutral, non-polar and aliphatic. These nine residues were designated as the core region in ICLIST-2 (Fig. 3B). While detailed analysis awaits structural determination via X-ray crystallography, these results suggest that the predictive model used in this study could explain the experimental results to a certain degree, enabling downstream protein engineering.

**Figure 3.**
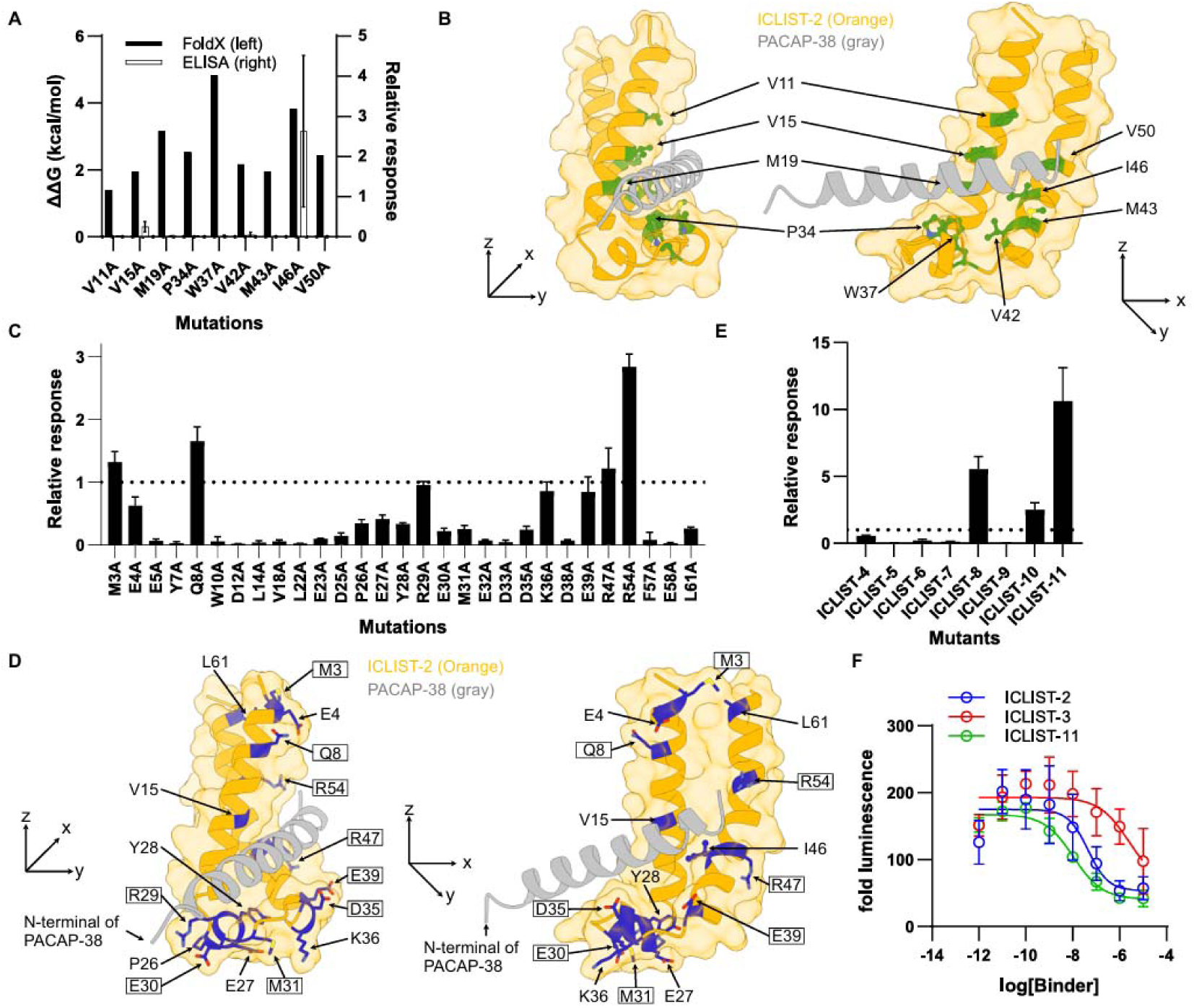
Design and evaluation of ICLIST-2 mutants. **A** Predicted ΔΔG (kcal/mol) values upon, using FoldX^30, 31^, and binding response measured by ELISA, for the nine alanine substitutions located in the vicinity of PACAP-38 (left axis). Binding response measured by ELISA (right axis). **B** Structural mapping (green) of the nine residues located in the vicinity of PACAP-38, which are defined as the core region. The side chains of these nine residues are shown as a ball-and-stick model in green, while ICLIST-2 is shown as an orange ribbon model and PACAP-38 as a grey ribbon model. **C** Alanine scanning of ICLIST-2. The binding of each mutant to PACAP-38 was evaluated using an ELISA and is shown as a relative value, with the binding response of ICLIST-2 set to 1; the dotted line represents this value. **D** Structural mapping (blue) of residues that retained at least 20% of their activity compared to ICLIST-2 in the ICLIST-2 alanine scanning experiment. The side chains of these residues are shown in blue as a ball-and-stick model, while ICLIST-2 is shown in orange as a ribbon model and PACAP-38 in grey. The residues enclosed in a box indicate those into which mutations were introduced in ICLIST-11. **E** The binding of eight variants (ICLIST-4 to ICLIST-11, designed using ProteinMPNN based on ICLIST-2 binding to PACAP-38) is shown as the binding response measured by ELISA. Values are expressed as relative values, with the binding response of ICLIST-2 set to 1, and the dotted line represents the relative binding response of ICLIST-2. All experiments were performed in triplicate, and the results are shown as the mean ± S.D. **F** Concentration-dependent inhibition of PACAP (1–38)-induced cAMP responses by binders (ICLIST-2, ICLIST-3, and ICLIST-11) measured by the GloSensor cAMP assay. Expi293F cells transiently co-expressing the Glo-22F cAMP biosensor and PAC1R were treated with 1 nM PACAP (1– 38) pre-incubated with serial dilutions of each binder. The data point plotted at log(-12) represents the response to PACAP (1–38) alone (without binder). Data were fitted using a four-parameter sigmoidal curve (Sigmoidal, 4PL, X is concentration). Data are presented as mean ± SEM (n = 3).

Therefore, based on this predictive model and to further identify residues outside the core region important for PACAP-38 recognition, we systematically substituted residues at the interface with alanine and evaluated binding via ELISA (Fig. 3C and Fig. S5). Interestingly, the R54A and Q8A mutants exhibited average responses approximately 2.8- and 1.4-fold higher than ICLIST-2, respectively. SPR analysis revealed *K*_D_ values of 16.3 and 18.7 nM, respectively, indicating increased affinity for PACAP-38 (Fig. S6A, B and Table S3). These residues are both located near the C-terminal region of PACAP-38 (Fig. 3D), suggesting that optimising them could further increase the affinity of ICLIST-2.

### Improvement of PACAP-38 affinity of ICLIST-2

Although ICLIST-2 exhibited a relatively high binding affinity for PACAP-38, we investigated the potential for further affinity enhancement through protein engineering. Based on the results of the alanine scanning experiment (Fig. 3C), we hypothesized that residues for which binding affinity was not lost upon Ala substitution could be further optimized. We initially focused on six residues where activity was enhanced or maintained (M3A, E4A, E39A, I46A, R47A and R54A). We prepared a combinatorial library using NNK primers and screened it via phage display. However, we were unable to obtain any mutants that outperformed ICLIST-2 (data not shown). This finding emphasizes that the six mutations used in the combinatorial library based on ICLIST-2 were insufficient and highlights the need to search for variants within a broader search space. To verify the existence of ICLIST-2 variants with higher affinity within a broader search space, we turned to ProteinMPNN, a widely used tool for deep learning-based inverse folding (Fig. 1). Based on Fig. 3C, we selected 17 residues that maintained at least 20 per cent of the mean response of the control (highlighted in blue in Fig. 3D). All amino acid residues in ICLIST-2 other than these 17 were fixed and residue optimization was performed using ProteinMPNN. As Bindcraft lacks information on the 11 C-terminal residues of PACAP-38, correctly evaluating interactions with this disordered region is necessary to improve affinity for PACAP-38. Therefore, complex models of full-length PACAP-38 and ICLIST-2 were generated using AlphaFold3^32^ and Boltz-2^33^ as the input for ProteinMPNN^34^. The AlphaFold3 prediction produced a model in which the disordered region was represented as a coil. However, due to the high flexibility of the 11 C-terminal residues, the arrangement differed across the five output models. Consequently, all five top-ranked models were provided to ProteinMPNN. Unlike the AlphaFold3 predictions, the Boltz-2 predictions represented the disordered region as an α-helix and two distinct conformations were observed (see Fig. S7). The pLDDT scores were low for the coils predicted by AlphaFold3, while those predicted by Boltz-2 were generally higher (Fig. S7). At the design stage, neither possibility was ruled out and 50 sequences were independently generated for each of the seven model structures using ProteinMPNN. Of these 350 sequences, eight were selected through *in silico* screening using two criteria: confidence score ≥ 0.940 and iPTM ≥ 0.954 (Fig. S8 and Table S4). When the binding of these variants to PACAP-38 was evaluated by ELISA, three variants exhibited a higher response than ICLIST-2 (Fig. 3E). All three of these variants were designed using AlphaFold3 models as input; the models used for ICLIST-4, 8, and 11 were different from one another. This suggests that AlphaFold3 is a suitable method for predicting the ICLIST-2–PACAP-38 complex model when designing ICLIST-2 variants using ProteinMPNN. In the ELISA screen, ICLIST-8 and ICLIST-11 exhibited particularly high binding responses, both showing an E_max_ more than five times higher than that of ICLIST-2 (Table 2). Interestingly, while ICLIST-8 showed a reduced apparent potency (higher EC_50_), ICLIST-11 maintained an EC_50_ comparable to ICLIST-2 despite this markedly elevated maximum response, highlighting ICLIST-11 as the most promising candidate (Fig. S9). Indeed, when the binding affinity of ICLIST-11 to PACAP-38 was quantitatively evaluated by SPR, the *K*_D_ was determined to be 4.51 nM, confirming a substantial improvement compared to ICLIST-2 (Table 1 and Fig. S6C).

**Table 2.** EC_50_ and E_max_ of ICLISTs.

|  | EC<sub>50</sub><br>(nM) | E<sub>max</sub><br>(% of ICLIST-2) |
|---|---|---|
| ICLIST-2 | 258.8 | 100 |
| ICLIST-8 | 941.9 | 506 |
| ICLIST-11 | 263.6 | 530 |

The binding rate (*k*_a_) increased by approximately 2.4-fold, while the dissociation rate (*k*_d_) decreased by 40 per cent. It is therefore considered that the nine mutations introduced into ICLIST-11 by ProteinMPNN (enclosed in boxes in Fig. 3D) improved both the binding and dissociation rates with PACAP-38. This resulted in an approximately 5.9-fold increase in affinity compared to ICLIST-2. To investigate this factor, the interactions between ICLISTs and PACAP-38 were predicted using PLIP calculations^35^. Focusing on the residues that differ between ICLIST-2 and ICLIST-11 revealed that ICLIST-11 acquired new hydrophobic interactions via M3V and R54W, along with a putative salt bridge mediated by Q8E (Tables S5 and S6). As mentioned earlier, a comparison of ICLIST-2 and ICLIST-3 suggested that Q8 might be important in the highly flexible region at the C-terminus of PACAP-38 (Fig. S3). The likelihood of establishing this additional electrostatic interaction through the Q8E substitution in ICLIST-11 potentially corroborates this conclusion. ICLIST-11 had fewer hydrophobic interactions, hydrogen bonds and π-stacking than ICLIST-2, while the number of salt bridges increased. This is thought to be due to slight changes in the binding mode with PACAP-38 caused by the nine mutations. The above analysis suggests that M3V, Q8E and R54W play important roles in ICLIST-11’s high binding affinity for PACAP-38.

Finally, we assessed whether the engineered ICLIST variants could inhibit PACAP-38-mediated activation of PAC1R in cell-based functional assays. Using a GloSensor cAMP assay in PAC1R-expressing Expi293F cells, the neutralizing activities of ICLIST-2, ICLIST-3, and the affinity-matured variant ICLIST-11 were evaluated against 1 nM PACAP-38-induced intracellular cAMP accumulation (Fig. 3F). Consistent with their binding affinities, all tested binders exhibited concentration-dependent inhibition of PACAP-38-induced responses. In particular, ICLIST-11 exhibited the highest inhibitory potency, shifting the curve significantly toward lower concentrations with an IC_50_ of 10.6 nM, which represents an approximately 4.3-fold improvement over parental ICLIST-2 (IC_50_ = 45.0 nM) and substantially outperforms ICLIST-3 (IC_50_ = 2.5 μM). Furthermore, while ICLIST-3 failed to achieve complete inhibition within the tested concentration range, both ICLIST-11 and ICLIST-2 efficiently suppressed the luminescence signal back toward basal levels at micromolar concentrations. These results demonstrate that the enhanced binding affinity of ICLIST-11 successfully translates into superior neutralizing efficacy and potency against PACAP-38-induced PAC1R activation.

### Thermostability of ICLISTs

Next, we assessed the changes to the structure and physical properties of ICLIST-11. Like ICLIST-2, the CD spectrum of ICLIST-11 exhibited the typical characteristics of an α-helix, and secondary structure prediction indicated that the entire structure consisted of α-helices. Interestingly, when the thermal stability of the secondary structure was measured by CD, the T_m_ values were found to be 66.4 °C and 80.4 °C for ICLIST-2 and ICLIST-11, respectively. Thus, ICLIST-11 exhibited a 14 °C improvement in structural stability (Fig. 4A). Comparing the structural models of ICLIST-2 and ICLIST-11 revealed an RMSD of 0.805 Å for Cα atoms, indicating no significant differences in the overall structure (Fig. 4B). Furthermore, although nine amino acid substitutions were introduced in ICLIST-11, analysis using ChimeraX revealed no new hydrogen bonds resulting from these mutations (Fig. S10).

**Fig. 4.**
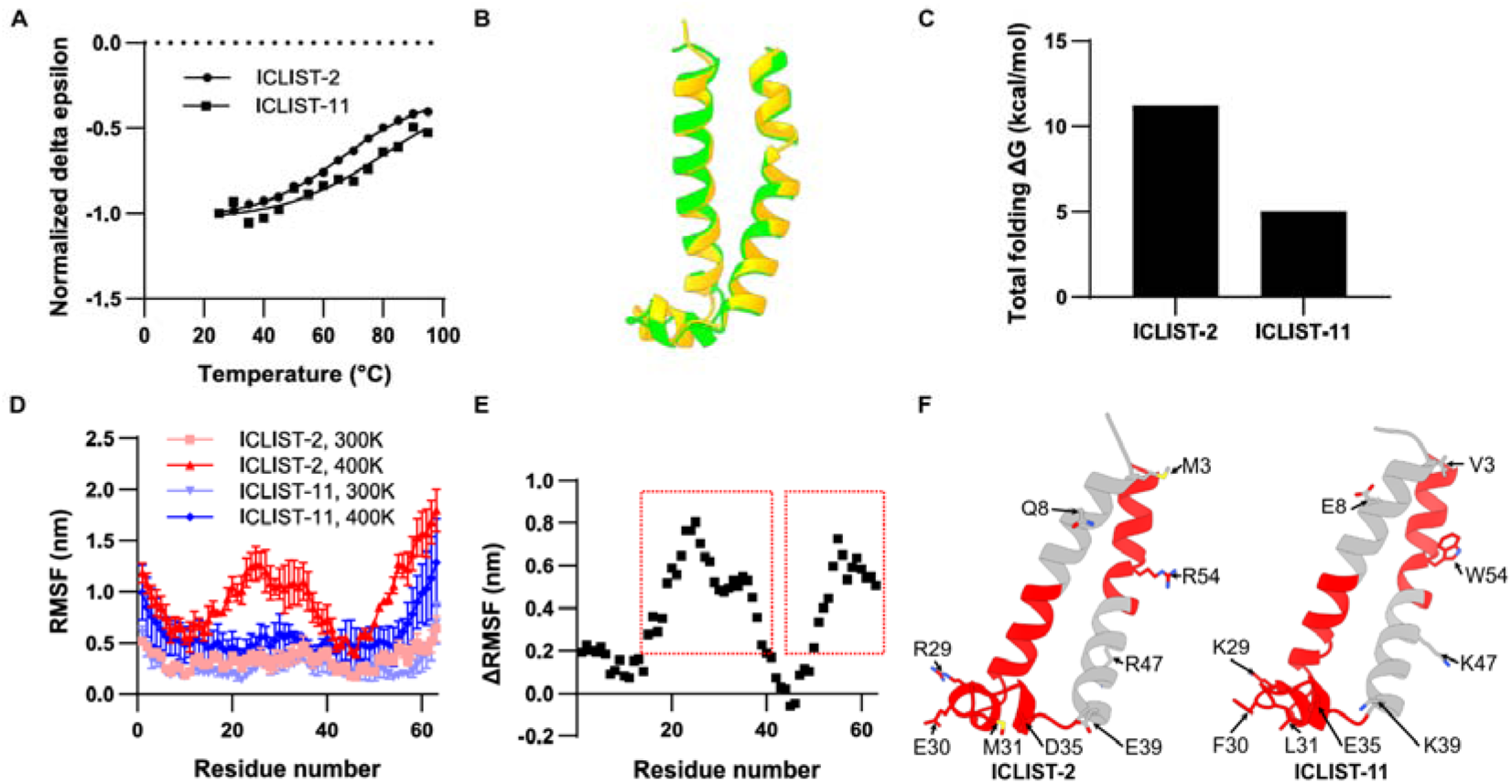
Thermostability and MD simulation of ICLISTs. **A** The temperature-dependent variation in the 220 nm value in the CD spectra of ICLIST-2 and ICLIST-11. The T_m_ values, as calculated using GraphPad Prism 10.6.1, were found to be 66.4 °C and 80.4 °C for ICLIST-2 and ICLIST-11, respectively. **B** Superimposition of the overall structures of ICLIST-2 (yellow) and ICLIST-11 (green). The RMSD for Cα atoms was 0.805 Å. **C** The ΔΔG (kcal/mol) values for ICLIST-2 and ICLIST-11 were calculated using FoldX^30, 31^. **D** RMSF (nm) obtained from MD simulations of ICLIST-2 (red) and ICLIST-11 (blue). MD simulations were performed three times at 300 K and 400 K, respectively. **E** Shows the difference in RMSF values for each residue, calculated at 300 K and 400 K. The region enclosed by the red dotted line indicates areas where the difference in RMSF was 0.25 nm or greater. **F** Ribbon models of the three-dimensional structures of ICLIST-2 (left) and ICLIST-11 (right), showing regions where a difference of 0.25 nm or more was observed in the RMSF calculated from the 300 K and 400 K MD simulations. The side chains of the nine residues that differ between ICLIST-2 and ICLIST-11 are displayed as stick models.

Conversely, energy calculations performed with FoldX software^30, 31^ indicated that ICLIST-11 exhibited a lower folding free energy (ΔG), corresponding to a negative ΔΔG ralative to ICLIST-2 and suggesting structural stabilisation (Fig. 4C). To examine this mechanism in greater detail, molecular dynamics simulations were carried out. While no significant differences in RMSF were observed in simulations at 300 K, calculations at 400 K revealed a substantial increase in the RMSF value for ICLIST-2 at residue number 30 (Fig. 4D, E). Structural mapping of this region revealed that ICLIST-11 introduced three hydrophobic residues (E30F, M31L and R54W) (Fig. 4F). This suggests that the structure of ICLIST-11 is stabilised by hydrophobic interactions centred on these three residues. However, no clear difference in RMSD over time was observed between ICLIST-2 and ICLIST-11 (Fig. S11). This indicates that, although ICLIST-2 and ICLIST-11 have virtually the same overall structure, the introduction of mutations in hydrophobic residues has made the ICLIST-11 structure more rigid. It is therefore likely that optimisation using ProteinMPNN allowed ICLIST-11 to retain the overall structure of ICLIST-2 whilst improving the thermal stability of the entire molecule by acquiring hydrophobic interactions resulting from the E30F, M31L, and R54W mutations.

### Plasma assays for ICLIST reactivity

To function as a therapeutic, ICLIST constructs would need to be compatible with functioning in the blood of patients. We therefore examined the reactivity of ICLIST-2 and ICLIST-11 (Fig. 5A). No reaction was observed in the presence of serum albumin, the main protein in blood plasma. Furthermore, despite concerns about non-specific electrostatic interactions, due to the large number of negatively charged glutamic acid residues in the ICLISTs (Table S1 and S4), there was almost no reaction with poly-D-lysine. These results suggest that the ICLISTs exhibit low non-specific background binding to abundant plasma proteins and charged polymers. Next, the stability of ICLISTs in blood was evaluated. No band fragmentation was observed on SDS-PAGE when ICLIST-2 and ICLIST-11 were stored in plasma at 37 °C for 24 hours (Fig. 5B and S12). Furthermore, after 24 hours of storage, when the ICLISTs were returned to buffer and their binding to PACAP-38 was assessed, it was found to be 70% of that observed in the control (Fig. 5C). This suggests that, although the peptide bonds of the ICLISTs were not cleaved by proteases in plasma, their binding capacity to PACAP-38 may have been reduced by compounds inhibiting their binding or by chemical changes in the ICLISTs themselves. Conversely, when the binding capacity in plasma was evaluated at concentrations ranging from 0% to 100%, a significant decrease in the binding response was observed in proportion to the plasma concentration (Fig. 5D,E). This phenomenon was more pronounced in ICLIST-2, suggesting that ICLIST-11 is more resistant to plasma components than ICLIST-2. In particular, when the plasma concentration reached 100%, the ligand-binding signal decreased to 10–20% compared to the buffer-only condition. However, this was not due to the binder being inactivated, as the binding capacity remained at 70% when measured after the sample had been left in plasma for one day and then returned to buffer. Consequently, this reduction in signal is not attributed to irreversible denaturation or degradation of the molecule, but rather to a foreign substance at high concentration (e.g. 100% plasma containing background proteins). These substances could be physically inhibiting the detection of ligand binding in the assay system (e.g. ELISA or SPR) (matrix effect), or leading to reversible masking. Based on the above evaluation, it is clear that, while ICLISTs are resistant to cleavage by proteases in mouse plasma, their binding to PACAP-38 may be inhibited due to the matrix effect. Taking these factors into account, future animal experiments must elucidate the actual *in vivo* binding behaviour of ICLISTs, and further research is anticipated.

**Figure 5.**
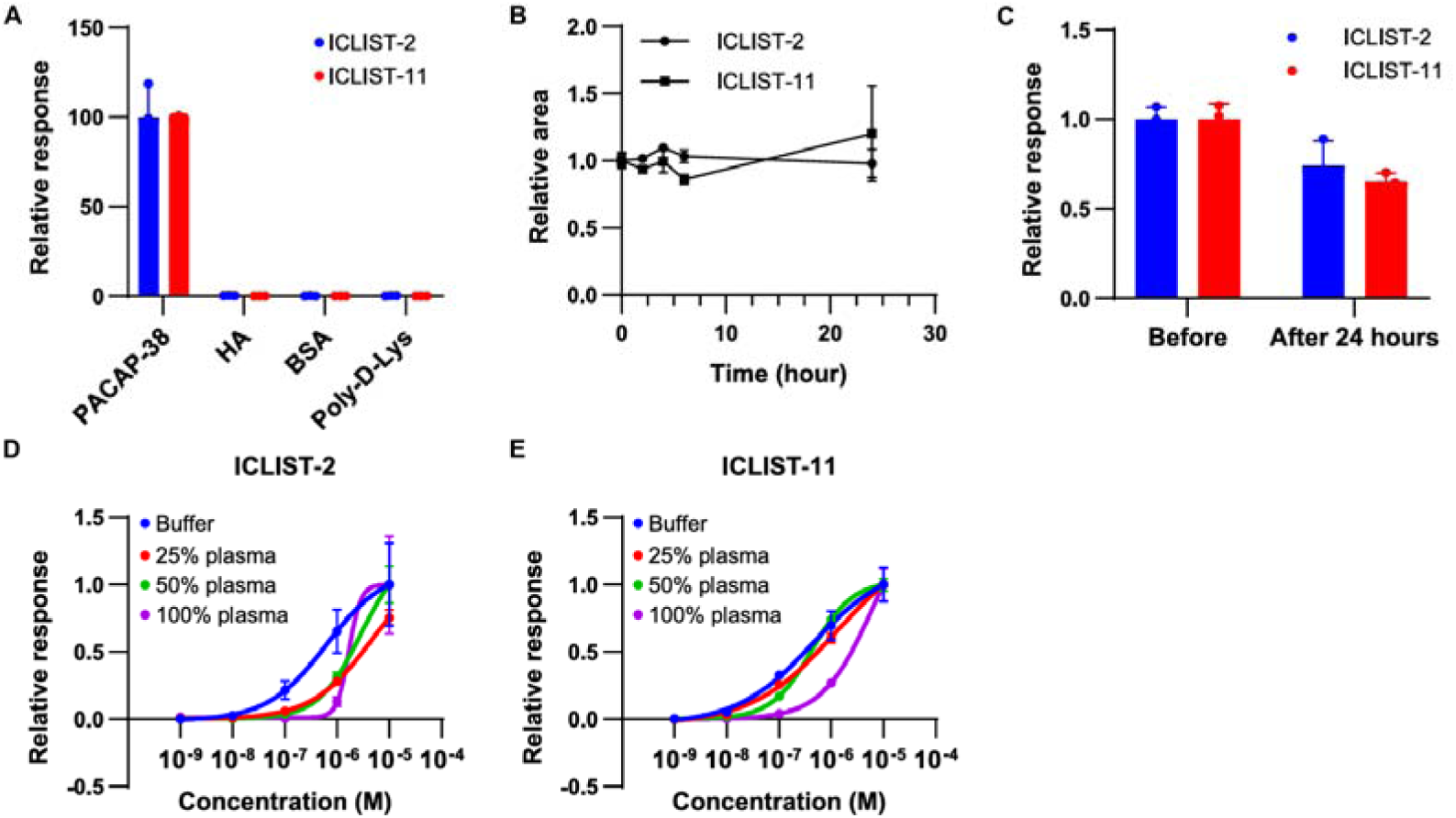
Mouse plasma assay. **A** Selectivity of ICLISTs. HA; Human albumin and BSA; bovine albumin. **B** Storage stability of ICLIST-2 and ICLIST-11 in mouse plasma, shown as the relative SDS-PAGE band area quantified by ImageJ at 0, 1, 3, 6, and 24 hours relative to a baseline of 1 before storage. **C** Binding capacity to PACAP-38 before and after 24 hours’ storage in mouse plasma. **D,E** Binding response to PACAP-38 in 0, 25, 50 and 100% plasma. ICLIST-2 **D** and ICLIST-11 **E** are shown respectively. All experiments were performed in triplicate; the mean ± S.D. is shown.

## Conclusion

A mini-protein that targets PACAP-38 was designed using Bindcraft, and three binders that exhibit binding affinity were identified. Of these, ICLIST-2 exhibited particularly high affinity with a *K_D_* of 26.5 nM. To further improve affinity, 17 different residues were selected for optimisation using ProteinMPNN, based on information from alanine scanning. Consequently, ICLIST-11, which incorporated nine mutations into ICLIST-2, exhibited the highest affinity with a *K_D_* of 4.51 nM. The acquisition of novel interactions with PACAP-38 via M3V, Q8E and R54W was suggested to play a significant role. Furthermore, ICLIST-11 had a T_m_ 14 °C higher than ICLIST-2 due to the formation of hydrophobic interactions via E30F, M31L and R54W. Moreover, cell-based functional assays demonstrated that ICLIST-11 effectively neutralized PACAP-38-induced PAC1R activation in a concentration-dependent manner, exhibiting markedly superior inhibitory potency and efficacy compared to parental ICLIST-2 (IC50 = 10.6 nM). These findings provide important foundational information for designing PACAP-38 binders, which act as ligand traps function in drug discovery.

## Materials and methods

### Strains and media

Standard DNA cloning was performed with chemically competent DH10B and BL21 (DE3) (Thermo Fisher Scientific). Cells were grown in Luria-Bertani medium (Sigma-Aldrich), 2 × TY medium (5 g / L NaCl, 10 g / L yeast extract, 16 g / L tryptone) or S.O.C. medium (Sigma-Aldrich). Kanamycin (50 μg / mL) and carbenicillin (100 μg / mL) were added where appropriate.

### Structure prediction

The structure models were generated using the Boltz-2^33^ and AlphaFold Server^32^ (AlphaFold 3 model, Google DeepMind) and visualized using ChimeraX version 1.10.1^36^.

### *De novo* binder generation and screening for PACAP-38

Target-specific binders were computationally designed *de novo* using the Bindcraft pipeline. All computational design trajectories, sequence optimizations, and structural simulations were executed utilizing the cloud-based high-performance computing platform provided by Ariax Bio. The structure of PACAP-38 was used as an initial template, based on the cryo-electron microscopy structure of the complex between the target protein, the human PAC1 receptor, and PACAP-38 (PDB ID: 6M1I). As the eleven residues from the 28th position were disordered, the structure from the N-terminus to the 27th position was used as the query in this design. To maximize structural prediction quality for challenging epitopes, the pipeline was configured with advanced settings utilizing the default 4-stage multimer (Hard target) protocol. Initial binder scaffolds were generated using the ColabDesign framework embedded within Bindcraft, with the binder length strictly constrained between 31 and 100 amino acids. A total of 209 computational design trajectories were executed on the Ariax Bio platform.

### Sequence Optimization and Filtering

For each generated backbone, sequence optimization was performed using ProteinMPNN to maximize interface complementarity and autonomous folding stability. To screen the generated designs, a multi-parametric scoring and filtering pipeline was applied utilizing the standard default parameters specified in the Bindcraft framework (default_filters.json). Rather than relying on PyRosetta, the physical and thermodynamic properties of the interface were evaluated using FreeBindCraft (https://github.com/cytokineking/FreeBindCraft), an open-source scoring alternative. Structural selection and filtering were strictly guided by the following default cutoffs: Binder Autonomous Folding Quality: Verified using a Predicted Local Distance Difference Test threshold (pLDDT≥80). Interface Structural Rigidity: Confirmed by a cross-interface Predicted Alignment Error threshold (PAE≤5.0). Interface Affinity and Packing: Constrained by the open-source predicted interface energy (ΔG), contact surface area, and shape complementarity metrics predefined in the standard filtering protocol.

### Structural Refinement and Final Selection

The candidate complexes that successfully cleared the initial filters underwent structural relaxation using the AMBER force field (via OpenMM) to eliminate steric clashes and optimize side-chain packing. Final verification was performed by re-predicting the complex structures using AlphaFold2-multimer without templates to confirm structural convergence. Designs that exhibited high structural similarity to the generation models (Interface RMSD≤1.5 Å) were prioritized. Through this computational screening pipeline, the trajectory execution achieved a structural generation success rate of 47.8%. This yielded a final library of exactly 100 top-ranking binders (out of 209 total trajectories) that successfully cleared all strict computational criteria and were selected for downstream experimental validation.

### Expression and Purification

The pET-26b plasmid in which each putative binder was cloned was constructed by Gibson assembly. The target proteins were fused with the TEV protease cleavage site, the Flag-tag and His-tag sequence (ENLYFQGGGSGGGGSTSDYKDDDDKDYKDDDDKDYKDDDDKGGSEHHHHHH) at the C-terminus. The plasmid was introduced into *E. coli* BL21 (DE3), and colonies were inoculated in LB medium containing 50 μg/mL kanamycin at 37°C. Affinity tag fusion proteins were expressed by induction using 0.1 mM isopropyl-β-D-thiogalactopyranoside when the OD_600_ reached 0.6–1.0. The temperature was decreased to 30 °C, followed by incubation for 16 h. Harvested cells were washed with saline, then re-suspended in B-PER containing DNase I and Lysozyme (Thermo Fisher Scientific). After incubation for 15 min at room temperature, the lysate was centrifuged for 5 min at 15,000 × g. The supernatant was added to a His SpinTrap TALON column (Cytiva), and bound proteins were washed with a wash buffer (50 mM Tris-HCl, 300 mM NaCl, 50 mM imidazole, pH 8.0) and eluted with an elution buffer (50 mM Tris-HCl, 300 mM NaCl, 250 mM imidazole, pH 8.0). The eluted fraction buffer was exchanged for a solution of 50 mM Tris-HCl and 100 mM NaCl at pH 8.0 using an Amicon 3K (Merck), after which the sample was immediately subjected to the following assays.

### Preparation of ICLIST-2 Mutants

Point mutations of ICLIST-2 were prepared by site-directed mutagenesis using KAPA HiFi DNA polymerase (Roche). Meanwhile, multiple mutants were synthesized as gene fragments by Twist Bioscience and cloned into the pET-26b(+) vector. The plasmids were introduced into *E. coli* DH10β for plasmid purification. Protein expression and purification followed the same procedure as shown above.

### Optimization using ProteinMPNN

Computational sequence optimization of the 63-residue protein ICLIST-2 was performed using ProteinMPNN^34^. The full-length three-dimensional structures of PACAP-38 and ICLIST-2, calculated using Boltz-2^33^ and AlphaFold3^32^, were used as input structures. Specifically, 17 target residues were selected for optimization, allowing 19 amino acid types as candidates (with the exclusion of cysteine), yielding a design library of 50 sequences per input. The 3D structures of all generated sequences were subsequently predicted using Boltz-2^33^. To identify high-confidence binders, an *in-silico* screening was conducted based on structural metrics. Mutants were filtered using strict thresholds of a confidence score≥0.940 and an iPTM≥0.954, successfully narrowing the library down to 8 candidate sequences for further evaluation.

### Enzyme-linked immunosorbent assay (ELISA)

The binding capacity of each binder was analyzed by ELISA, in triple repeats, using biotinylated PACAP-38 (HY-P2609F, MedChemExpress) as the antigen. The wells of 96-well immuno plate (436110, Thermo Fisher Scientific) were coated with 100 µl of 1 µg/mL streptavidin in ELISA Phosphate Coating Buffer (CB07100, Thermo Fisher Scientific), and incubated overnight at 4 °C. The plates were washed with washing buffer (25 mM Tris, 0.15 M NaCl, 0.05% Tween-20 at pH 7.5) (28352, Thermo Fisher Scientific) and blocked with blocking buffer (StartingBlock™ Blocking Buffer, 37538, Thermo Fisher Scientific) 60 min at 25 °C. After discarding the solution, 100 µL of 0.1 µg/mL biotinylated PACAP-38 in blocking buffer was applied and incubated at 25 °C for 60 min with shaking at 400 rpm, followed by 5-times washing with washing buffer. The plates were incubated for 60 min with 100 µL of lysate or purified protein in blocking buffer, washed with washing buffer 5-times, and incubated for another 60 min at 25 °C with shaking at 400 rpm, with peroxidase-conjugated mouse anti-FLAG tag antibody (MA1-91878-HRP, Thermo Fisher Scientific) diluted 1:4000 in blocking buffer. After another wash, the plates were developed with 100 µL of substrate solution (SuperSignal™ ELISA Pico Chemiluminescent Substrate, 37070, Thermo Fisher Scientific) for 5 min at room temperature. The luminescence intensity was measured using a plate reader (CLARIOstar Plus, BMG Labtech).

### Surface Plasmon Resonance (SPR)

Binding kinetics were determined using a Biacore S-200 system. Biotinylated PACAP-38 was dissolved in HBS-EP+ buffer (0.01 M HEPES, 0.15 M NaCl, 3 mM EDTA, 0.05% Surfactant P20, pH 7.6, Cytiva), and immobilized onto a SA sensor chip. The ICLIST analytes were adjusted to the desired concentration by serial dilution in HBS-EP+ buffer. The sensorgrams were obtained with an association time of 100 s, dissociation time of 300 s, and flow rate of 30 μL min^−1^. To determine the binding kinetics (*k*_a_, *k*_d_, and *K*_D_ values), the obtained sensor grams were analyzed by Biacore S200 Evaluation software v.1.0, using a 1:1 binding or bivalent analyte model.

### Circular dichroism (CD)

CD measurements were performed on a Chirascan spectrometer (Applied Photophysics, UK). Far-UV CD spectra for secondary structure analysis were recorded at 25 °C from 190 to 260 nm using a 1 mm path length quartz glass cuvette at a protein concentration of 0.5 mg/mL in 20 mM Tris-HCl, 100 mM NaCl pH 8.0. For thermal denaturation profiles, CD intensity was measured at 5°C intervals over a range of 25–95 °C. The baseline of the buffer was subtracted for all measurements. The melting temperature (*T*_m_) was calculated from the temperature dependence of the CD signal using the Graph Pad Prism 10.6.1.

### Prediction of structural stability

Structural stability and in silico alanine scanning mutagenesis were performed using the FoldX 4 software package^30, 31^. The overall structural stability (ΔG(kcal/mol)) of the ICLIST-2 was then evaluated. Subsequently, alanine scanning was executed using the AlaScan command, where target residues were sequentially mutated to alanine. The changes in binding affinity or stability (ΔΔG(kcal/mol)) upon mutation were calculated using the FoldX energy function.

### MD simulation

The molecular dynamics (MD) simulations were performed using the GROMACS package with the AMBER99SB-ILDN force field. The system was solvated in a Dodecahedron box of water, neutralized with 100 mM Na+/Cl- ions, and energy-minimized. Equilibration was carried out under NVT and NPT ensembles at 300 K or 400 K. Finally, a 100 ns production MD run was executed with a time step of 2 fs. Trajectory analysis (RMSD, RMSF) was carried out using Neurosnap, an online platform (Neurosnap Inc., available at: https://neurosnap.ai/Neurosnap).

### Mouse plasma analysis

Mouse plasma was obtained from Rockland as Non-Sterile In Sodium Heparin (D408-06-0050). Mouse plasma and ICLISTs were mixed in a 100:1 ratio and stored at 37 °C for one day; storage stability was then assessed by confirming the degradation of ICLISTs via SDS-PAGE. ImageJ was used to quantify the bands on the SDS-PAGE gel. The binding capacity in mouse plasma was evaluated using a solution prepared by mixing mouse plasma with ICLISTs at a fixed ratio, in accordance with the ELISA protocol described above.

### Cell-based assay

The inhibitory effects of the binders on PACAP-induced cAMP accumulation in PAC1R-expressing cells were measured using the GloSensor cAMP assay (Promega). Expi293F cells were transiently co-transfected with the Glo-22F cAMP biosensor-encoding plasmid and PAC1R plasmids using polyethyleneimine. After incubating the cells for 48–72 hours at 37 °C, the transfected cells were harvested and centrifuged at 190×g for 5 min, and resuspended in Hank’s balanced salt solution (HBSS, Thermo Fisher Scientific) containing 0.01% bovine serum albumin (BSA, Fujifilm Wako), 5 mM HEPES (pH 7.4), and 2 mM D-luciferin potassium solution (Fujifilm Wako). The cell suspension was seeded into a 96-well white plate (Corning) at 80 µL per well. PACAP (1-38) was prepared to give a final concentration of 1 nM and was mixed with a pre-prepared serial dilution of each binder before addition to the cells, and the mixture was incubated for 30 min at room temperature. The mixture was then added to the cells, which were incubated for 15 min at room temperature. The luminescence of the cells was measured using a SpectraMax i3x multi-plate reader. Data were analyzed using “Sigmoidal, 4PL, X is concentration” in GraphPad Prism 10.0.

## Data availability

All data required to evaluate the conclusions are included in the manuscript. Source data are provided with this paper.

## Acknowledgement

This work was supported by Ajinomoto Co., Inc. We are deeply grateful for the kind support provided by Kazutoshi Takahashi, Yasuaki Takakura, Masayuki Sugiki and Masaru Takayanagi (Ajinomoto Co., Inc.). We would like to thank Amir Guppy, Dario Bazzoli, Yuki Kimura, Olga Komarynets of Imperial College London for their valuable discussions. We would also like to thank Sam Horrel and Edward Bartlett of Imperial College London for their technological support.

## Author contributions

H.Y. and M.I. designed the research. H.Y. designed the proteins, screened them in silico, created the expression constructs, purified the proteins and performed ELISA, SPR, CD and MD simulation assays, as well as a mouse plasma assay. S.S. performed the cell-based assay. H.Y., S.S., Y.F. and M.I. wrote the manuscript. M.I. supervised the research. All authors read, revised and approved the manuscript.

## References

1. Collaborators GBDH. Global, regional, and national burden of migraine and tension-type headache, 1990-2016: a systematic analysis for the Global Burden of Disease Study 2016. Lancet Neurol 17, 954–976 (2018).

2. Stewart WF, Wood C, Reed ML, Roy J, Lipton RB, Group AA. Cumulative lifetime migraine incidence in women and men. Cephalalgia 28, 1170–1178 (2008).

3. Headache Classification Committee of the International Headache Society (IHS) The International Classification of Headache Disorders, 3rd edition. Cephalalgia 38, 1–211 (2018).

4. Ashina M. Migraine. New England Journal of Medicine 383, 1866–1876 (2020).

5. Moskowitz MA. The neurobiology of vascular head pain. Ann Neurol 16, 157–168 (1984).

6. Ashina M, Hansen JM, BO AD, Olesen J. Human models of migraine - short-term pain for long-term gain. Nat Rev Neurol 13, 713–724 (2017).

7. Ashina M, Hansen JM, Do TP, Melo-Carrillo A, Burstein R, Moskowitz MA. Migraine and the trigeminovascular system-40 years and counting. Lancet Neurol 18, 795–804 (2019).

8. Dodick DW. CGRP ligand and receptor monoclonal antibodies for migraine prevention: Evidence review and clinical implications. Cephalalgia 39, 445–458 (2019).

9. Khan S, Olesen A, Ashina M. CGRP, a target for preventive therapy in migraine and cluster headache: Systematic review of clinical data. Cephalalgia 39, 374–389 (2019).

10. Moreno-Ajona D, Perez-Rodriguez A, Goadsby PJ. Gepants, calcitonin-gene-related peptide receptor antagonists: what could be their role in migraine treatment? Curr Opin Neurol 33, 309–315 (2020).

11. Sacco S, et al. European headache federation guideline on the use of monoclonal antibodies acting on the calcitonin gene related peptide or its receptor for migraine prevention. J Headache Pain 20, 6 (2019).

12. Ernstsen C, et al. The PACAP pathway is independent of CGRP in mouse models of migraine: possible new drug target? Brain 145, 2450–2460 (2022).

13. Miyata A, et al. Isolation of a novel 38 residue-hypothalamic polypeptide which stimulates adenylate cyclase in pituitary cells. Biochem Biophys Res Commun 164, 567–574 (1989).

14. Al-Karagholi MA, Zhuang ZA, Beich S, Ashina H, Ashina M. PACAP38-induced migraine attacks are independent of CGRP signaling: a randomized controlled trial. J Headache Pain 26, 79 (2025).

15. Rawlings SR. PACAP, PACAP receptors, and intracellular signalling. Mol Cell Endocrinol 101, C5–9 (1994).

16. Miyata A, et al. Isolation of a neuropeptide corresponding to the N-terminal 27 residues of the pituitary adenylate cyclase activating polypeptide with 38 residues (PACAP38). Biochem Biophys Res Commun 170, 643–648 (1990).

17. Vollesen ALH, Amin FM, Ashina M. Targeted Pituitary Adenylate Cyclase-Activating Peptide Therapies for Migraine. Neurotherapeutics 15, 371–376 (2018).

18. Ashina M, et al. A phase 2, randomized, double-blind, placebo-controlled trial of AMG 301, a pituitary adenylate cyclase-activating polypeptide PAC1 receptor monoclonal antibody for migraine prevention. Cephalalgia 41, 33–44 (2021).

19. Ashina M, Phul R, Khodaie M, Löf E, Florea I. A Monoclonal Antibody to PACAP for Migraine Prevention. New England Journal of Medicine 391, 800–809 (2024).

20. Pellesi L, Ashina M, Martelletti P. Targeting the PACAP-38 pathway is an emerging therapeutic strategy for migraine prevention. Expert Opin Emerg Drugs 29, 57–64 (2024).

21. Lau JL, Dunn MK. Therapeutic peptides: Historical perspectives, current development trends, and future directions. Bioorg Med Chem 26, 2700–2707 (2018).

22. Muttenthaler M, King GF, Adams DJ, Alewood PF. Trends in peptide drug discovery. Nat Rev Drug Discov 20, 309–325 (2021).

23. Wang J, et al. Cryo-EM structures of PAC1 receptor reveal ligand binding mechanism. Cell Res 30, 436–445 (2020).

24. Chartrel N, Tonon MC, Vaudry H, Conlon JM. Primary structure of frog pituitary adenylate cyclase-activating polypeptide (PACAP) and effects of ovine PACAP on frog pituitary. Endocrinology 129, 3367–3371 (1991).

25. Hoyle CH. Neuropeptide families: evolutionary perspectives. Regul Pept 73, 1–33 (1998).

26. Sherwood NM, Krueckl SL, McRory JE. The origin and function of the pituitary adenylate cyclase-activating polypeptide (PACAP)/glucagon superfamily. Endocr Rev 21, 619–670 (2000).

27. Vaudry D, et al. Pituitary adenylate cyclase-activating polypeptide and its receptors: 20 years after the discovery. Pharmacol Rev 61, 283–357 (2009).

28. Segre GV, Goldring SR. Receptors for secretin, calcitonin, parathyroid hormone (PTH)/PTH-related peptide, vasoactive intestinal peptide, glucagonlike peptide 1, growth hormone-releasing hormone, and glucagon belong to a newly discovered G-protein-linked receptor family. Trends Endocrinol Metab 4, 309–314 (1993).

29. Pacesa M, et al. One-shot design of functional protein binders with BindCraft. Nature 646, 483–492 (2025).

30. Guerois R, Nielsen JE, Serrano L. Predicting changes in the stability of proteins and protein complexes: a study of more than 1000 mutations. J Mol Biol 320, 369–387 (2002).

31. Schymkowitz J, Borg J, Stricher F, Nys R, Rousseau F, Serrano L. The FoldX web server: an online force field. Nucleic Acids Res 33, W382–388 (2005).

32. Abramson J, et al. Accurate structure prediction of biomolecular interactions with AlphaFold 3. Nature 630, 493–500 (2024).

33. Passaro S, et al. Boltz-2: Towards Accurate and Efficient Binding Affinity Prediction. bioRxiv, (2025).

34. Dauparas J, et al. Robust deep learning–based protein sequence design using ProteinMPNN. Science 378, 49–56 (2022).

35. Adasme MF, et al. PLIP 2021: expanding the scope of the protein-ligand interaction profiler to DNA and RNA. Nucleic Acids Res 49, W530–W534 (2021).

36. Pettersen EF, et al. UCSF ChimeraX: Structure visualization for researchers, educators, and developers. Protein Science 30, 70–82 (2021).

