## Supplementary information for "*De novo* design of mini-proteins as specific binders to PACAP-38, a key mediator of migraine"

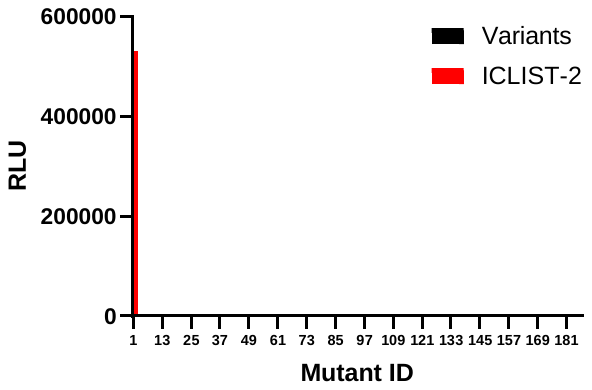

**Figure S1** **Screening of smaller binders for PACAP-38 except Top 20**

This figure shows the binding responses to PACAP-38 of 55 binders, ranging from 31 to 80 residues, excluding the top 20 designed using Bindcraft.

**A	B**

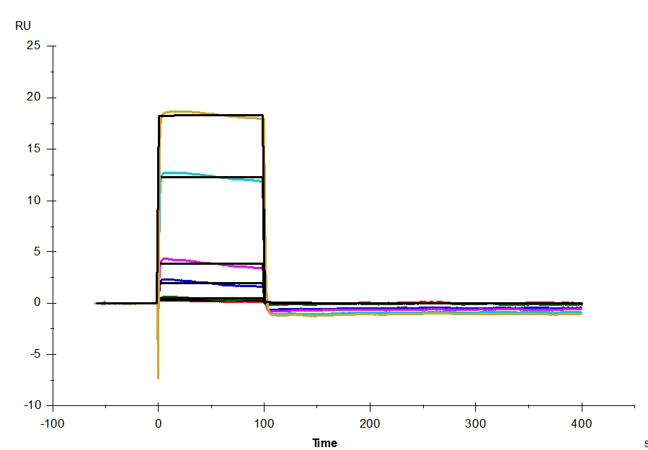

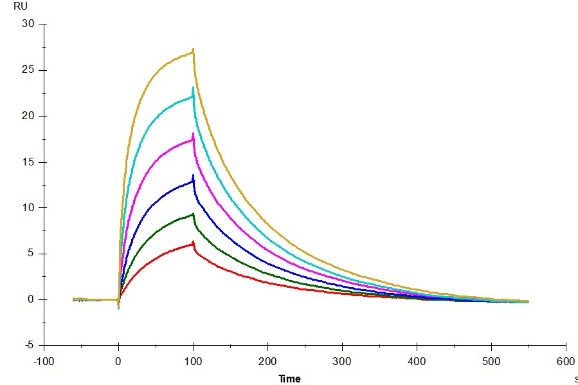

**C	D**

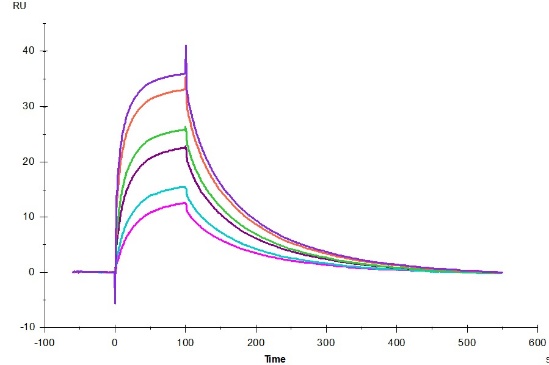
　
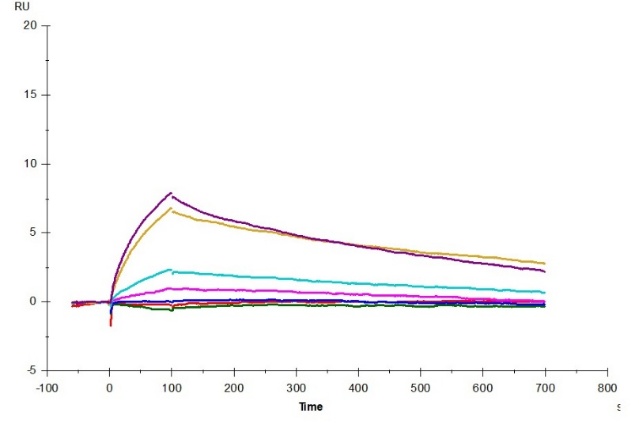

**Figure S2 Sensor grams of ICLISTs in surface plasmon resonance**

**A** ICLIST-1, **B** ICLIST-2, **C** ICLIST-3, **D** ICLIST-2 tandem

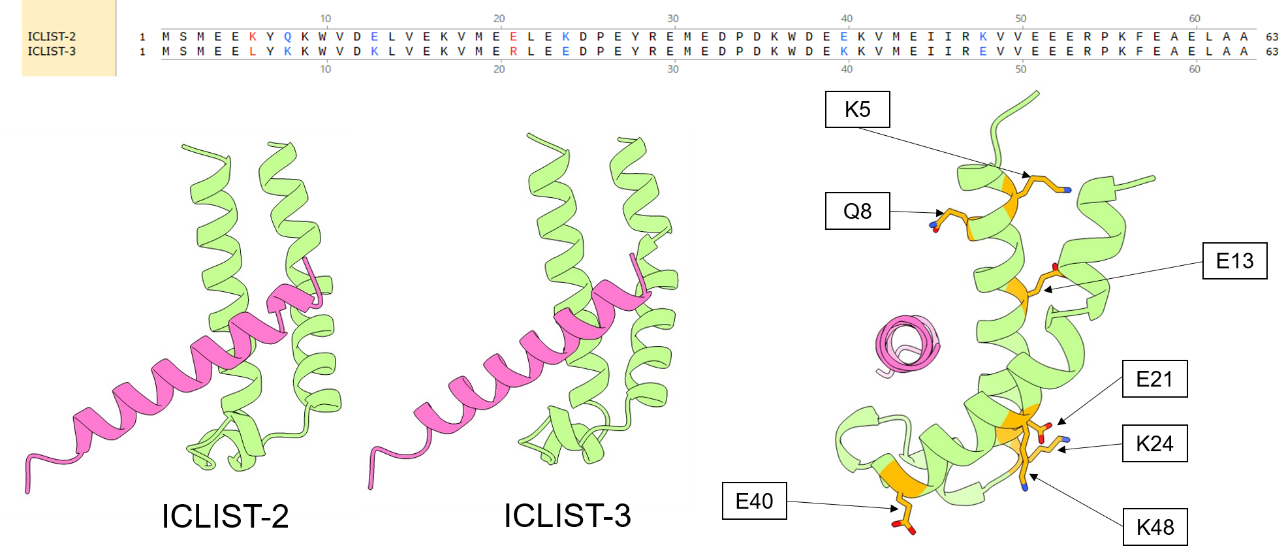

**Figure S3 Comparison between ICLIST-2 and ICLIST-3**

(upper) Sequence alignment, (lower left and center) Predicted structure of ICLIST-2 and ICLIST-3 with PACAP-38 (lower right) Structural mapping of the seven residues (marked in orange) that differ between ICLIST-2 and ICLIST-3 in ICLIST-2

**A	B**

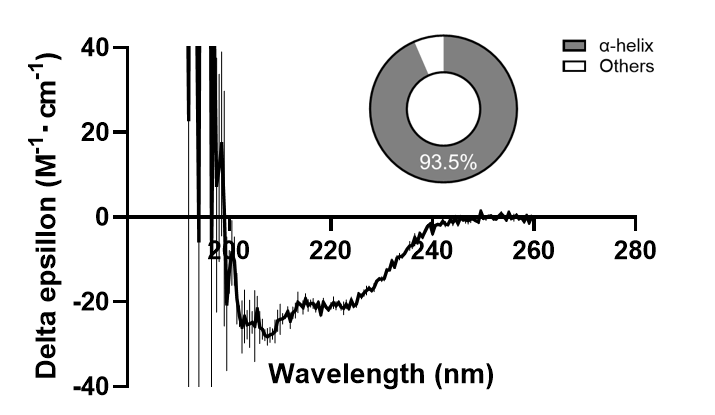

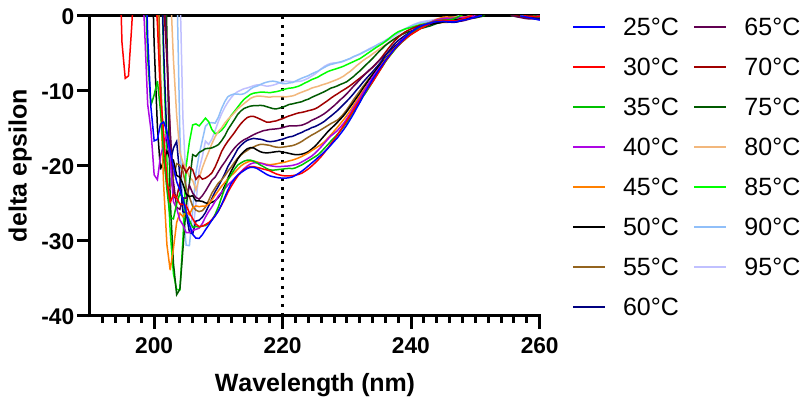

**C	D**

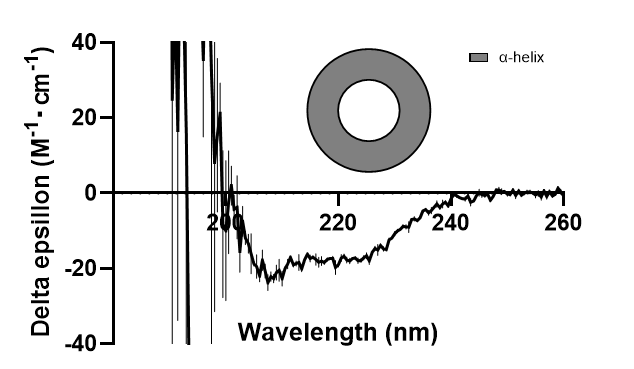

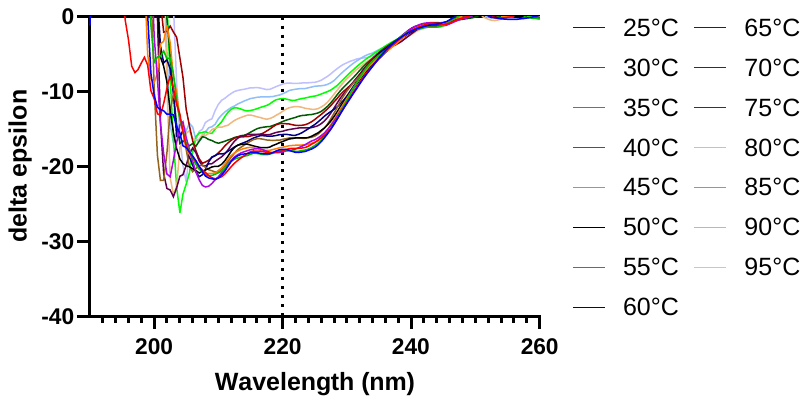

**Figure S4 CD spectrum of ICLIST-2 and ICLIST-11**

**A, C** CD spectra of ICLIST-2 (**A**) and ICLIST-11 (**C**). The pie charts show the proportions of secondary structures predicted by BeStSel for each. For ICLIST-2, 93.5% was classified as α-helix, with the remainder classified as the ‘Others’ component. Here, ‘Others’ mostly reflects irregular or unordered structures and is defined as a collection of all secondary structure elements excluded from the helix, β-sheet and turn components. **B, D** Changes in the CD spectra of ICLIST-2 (**B**) and ICLIST-11 (**D**) from 25°C to 95°C. The results, measured at 5°C intervals, are shown.

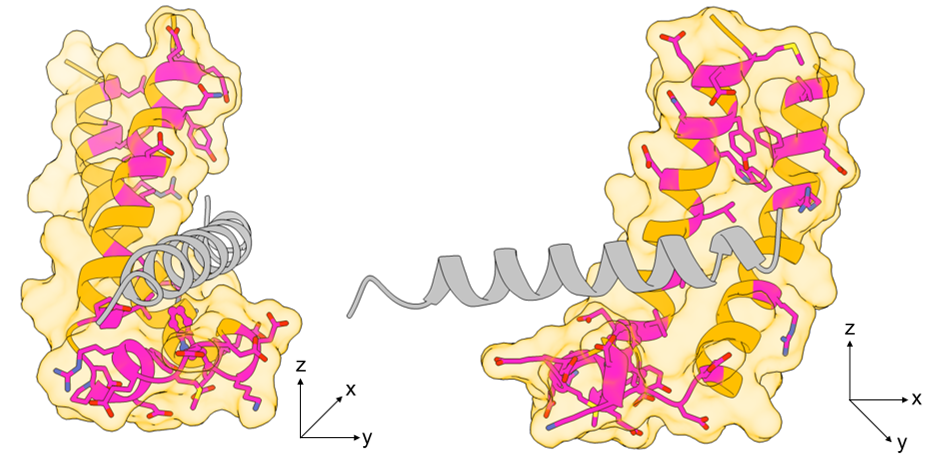

**Figure S5 Mutated residues in alanine scanning**

The structure of ICLIST-2 is shown using an orange ribbon model and a surface model; residues substituted with alanine in the alanine scanning experiment are shown in magenta, with their side chains depicted as a stick model. PACAP-38 is shown as a grey ribbon model.

**A	B**

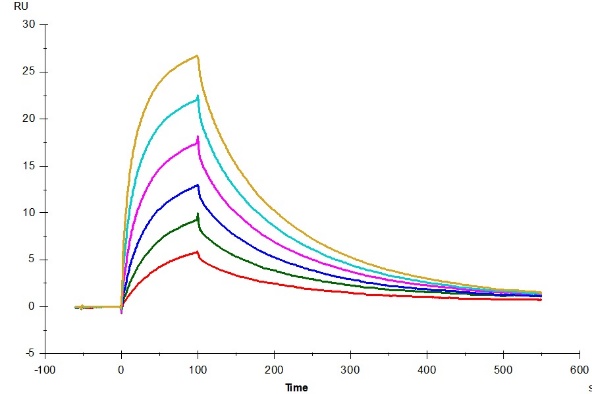

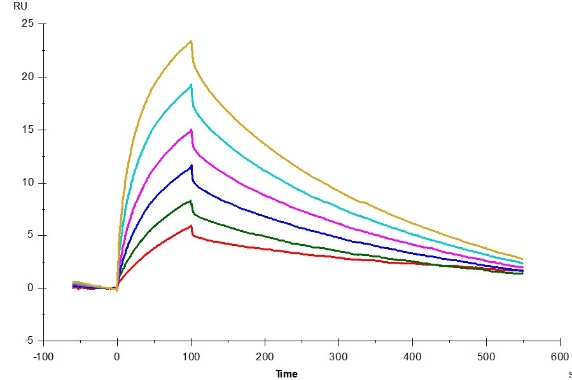

**C**

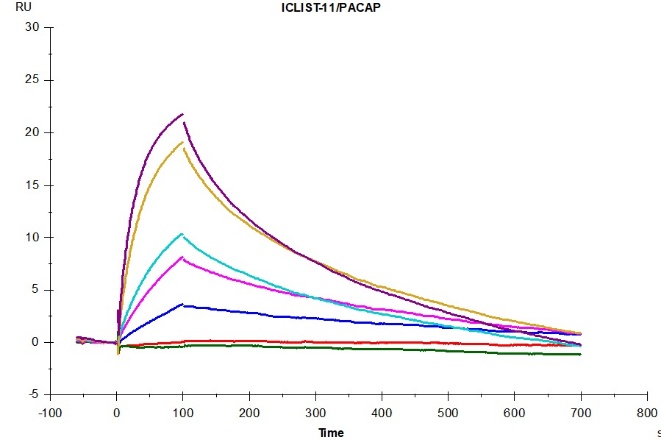

**Figure. S6 Sensorgrams of mutants of ICLIST-2 in surface plasmon resonance**

**A** ICLIST-2 (Q8A), **B** ICLIST-2 (R54A), **C** ICLIST-11

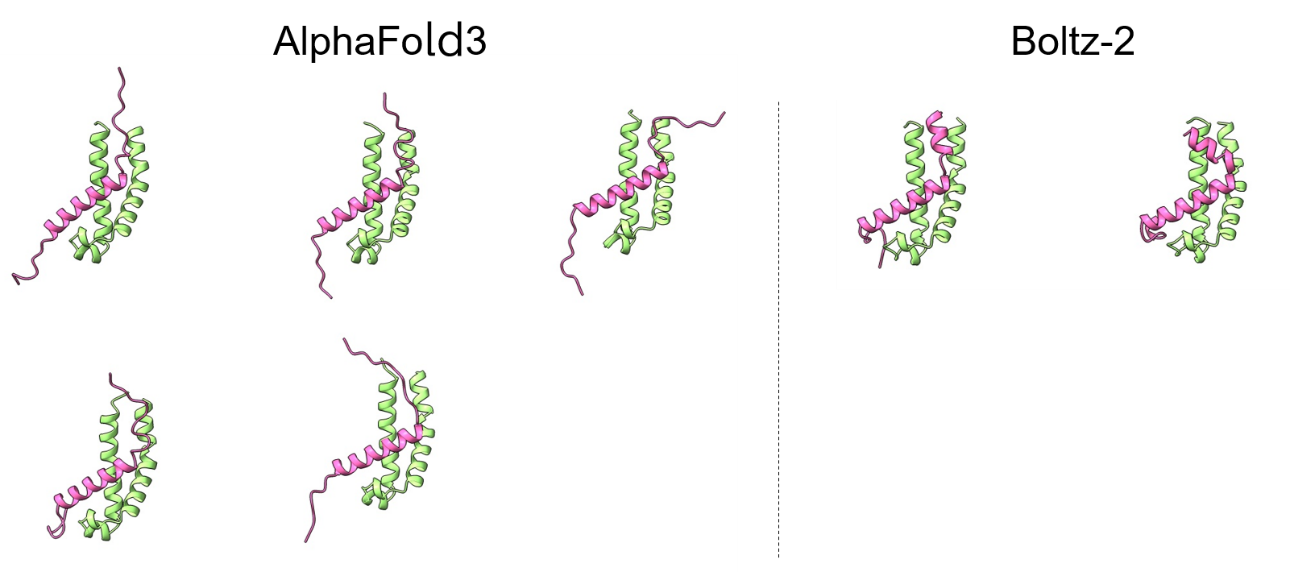

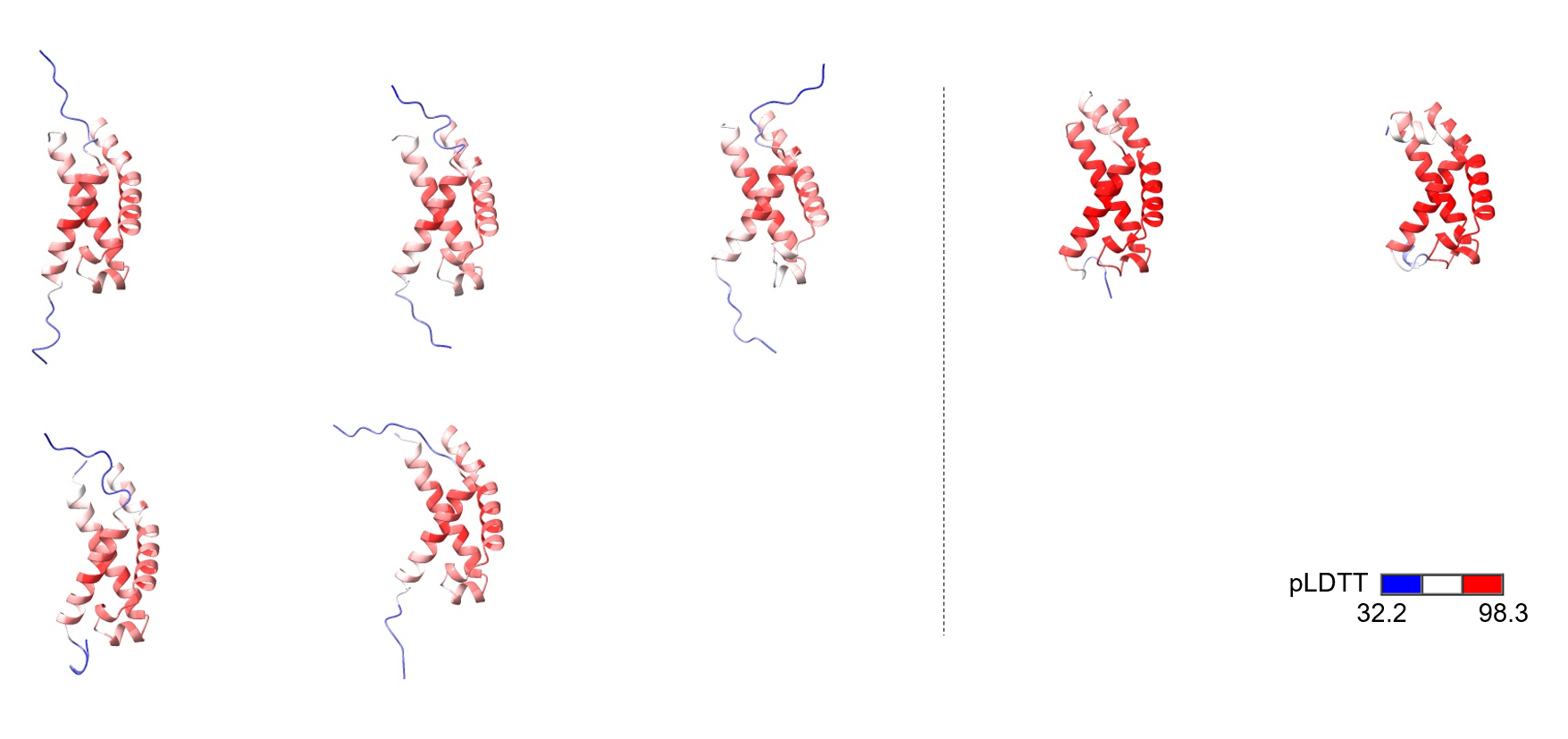

**Figure. S7 Structural prediction of the complex of ICLIST2 with full length of PACAP-38**

(upper left) Predicted models by AlphaFold3, (upper right) by Boltz-2 (lower left and right) Predicted models by AlphaFold3 and Boltz-2 colored by pLDTT

**A	B**

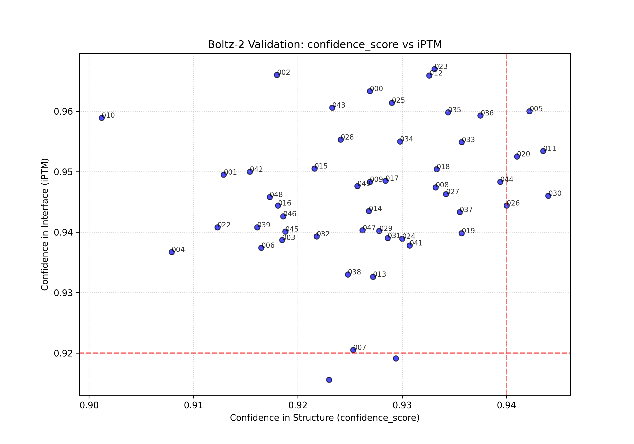

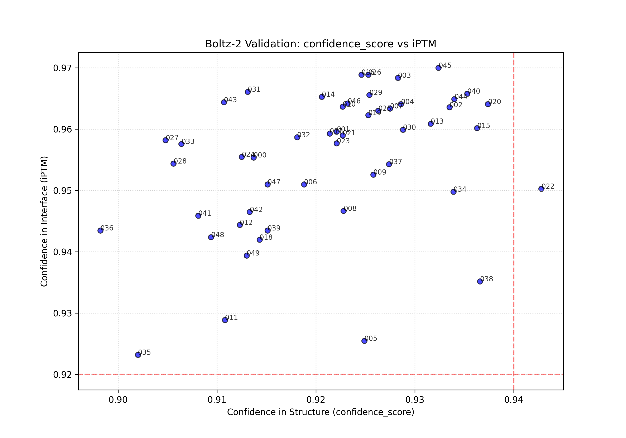

**C	D**

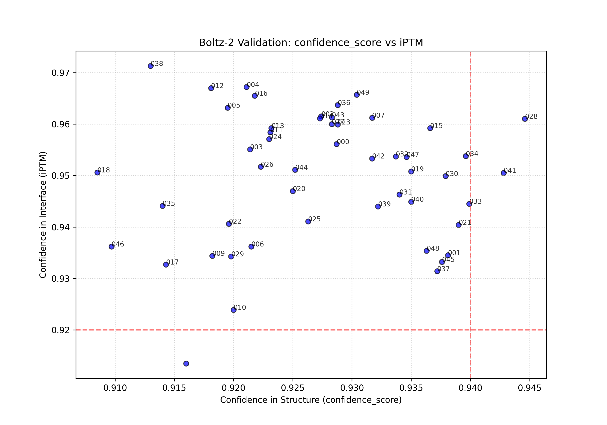

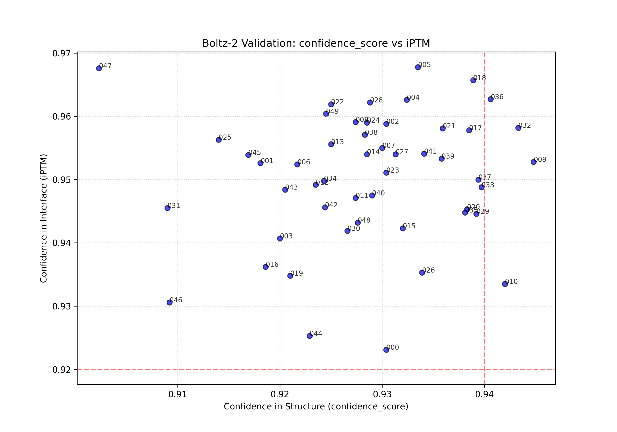

**E	F**

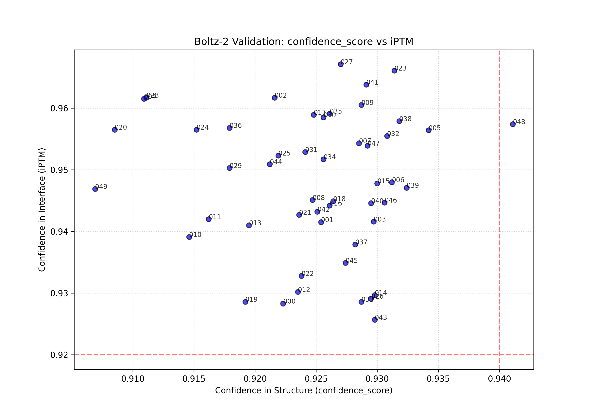

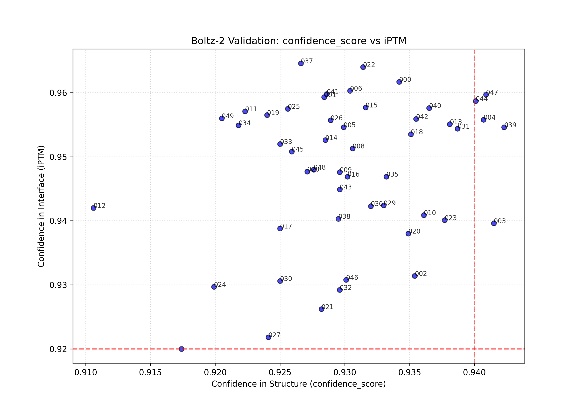

**G**

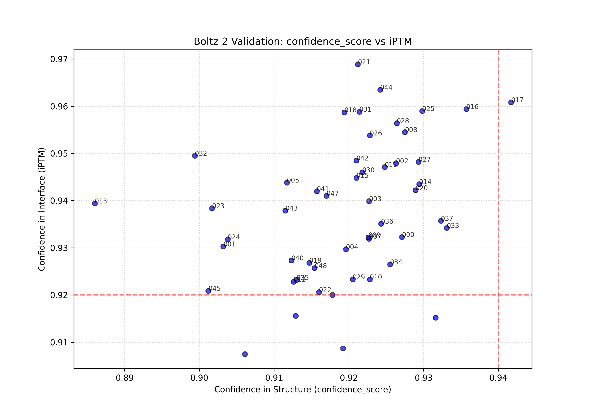

**Figure. S8 Plot of confidence scores and iPTM values in the predicted structural models of the ICLIST-2 and the full-length PACAP-38**

A plot of the confidence scores for the predicted structures against iPTM scores. **A, B, C, D** and **E** show 50 variants designed by ProteinMPNN based on structures predicted by AlphaFold3, whilst **F** and **G** show the results for two structures predicted by Boltz-2.

**A	B**

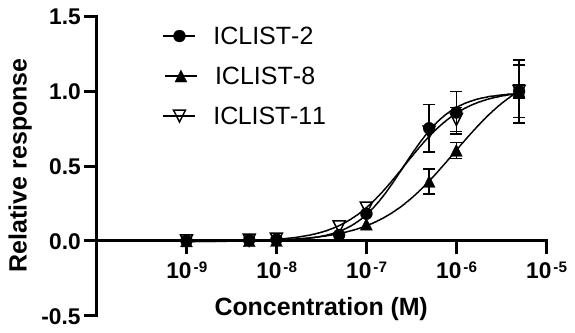

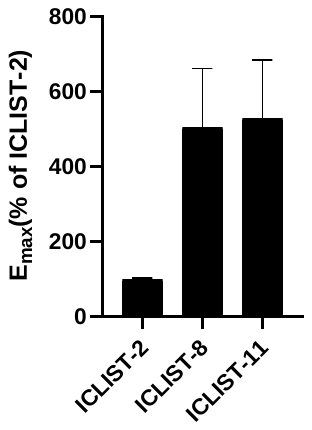

**Figure. S9 EC_50_ and Emax of mutants of ICLIST-2**

**A** The results of an ELISA assessment of the concentration-dependent binding responses of ICLIST-2, 8 and 11 to 0.1 μg/mL PACAP-38 are shown. **B** The relative Emax values for ICLIST-8 and ICLIST-11 are shown, with the Emax value for ICLIST-2 set at 100%.

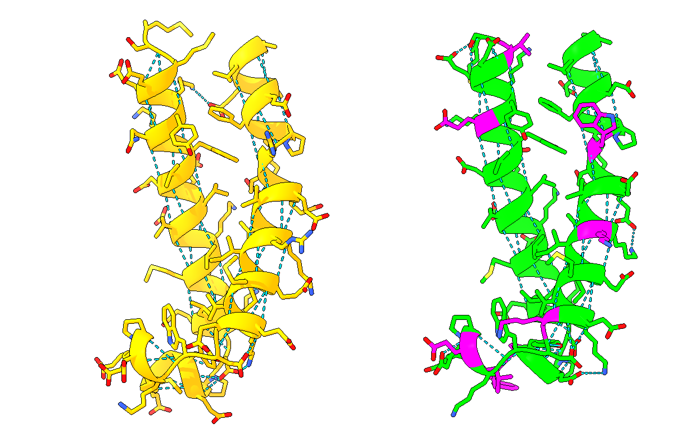

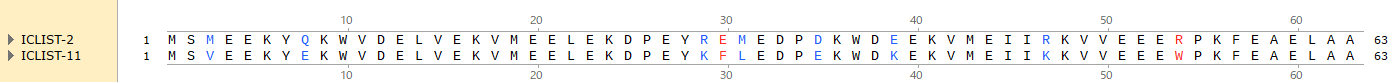

**Figure S10. Intramolecular hydrogen bonds of ICLIST-2 and ICLIST-11**

ICLIST-2 (yellow) and ICLIST-11 (green) are shown as ribbon models and hydrogen bonds are shown as dotted line. Magenta is 9 mutation which were introduced to ICLIST-11 from ICLIST-2.

**A	B**

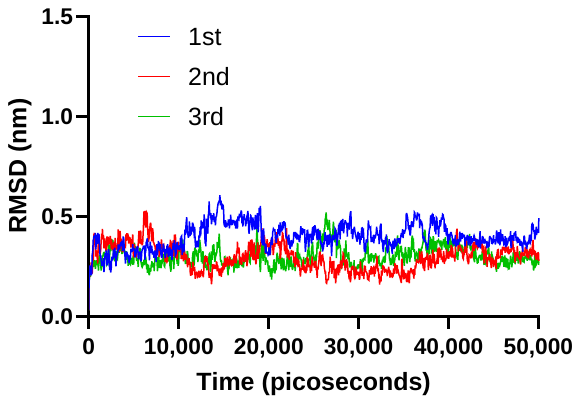

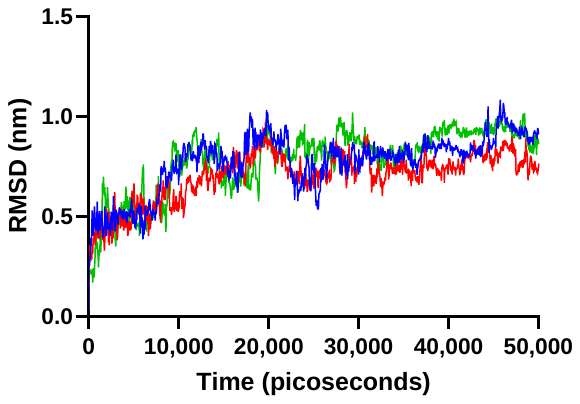

**C	D**

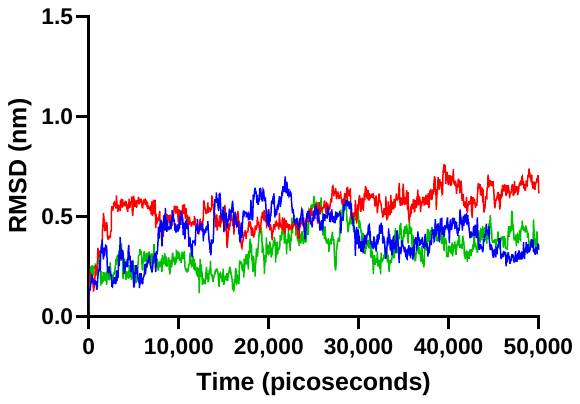

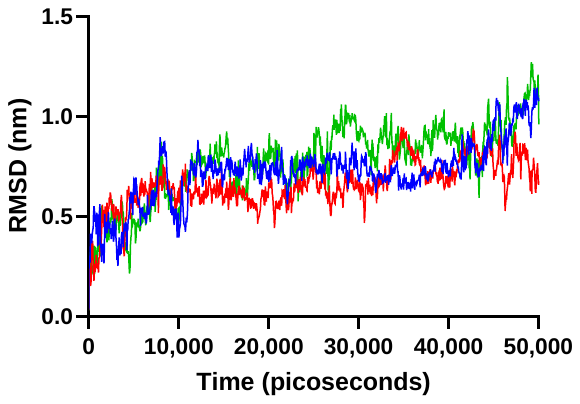

**Figure. S11 RMSD in MD simulation of ICLIST-2 and ICLIST-11**

**A, B** The RMSD for ICLIST-2, calculated three times at 300K and 400K. **C, D** The RMSD for ICLIST-11, calculated three times at 300K and 400K.

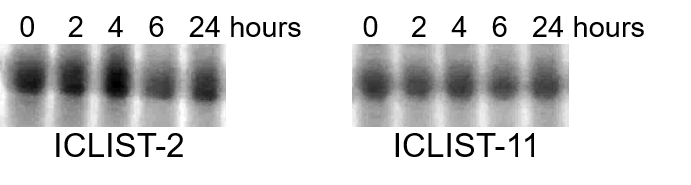

**Figure. S12 Storage stability of ICLIST-2 and ICLIST-11 in mouse plasma**

SDS-PAGE following incubation in mouse plasma at 37 °C for 0, 2, 4, 6 and 24 hours. The reaction in each sample was halted by the addition of SDS-PAGE sample buffer.

**Table S1 Amino acid sequence**

| ID | Sequence |
| --- | --- |
| PACAP-38 | HSDGIFTDSYSRYRKQMAVKKYLAAVLGKRYKQRVKNK |
| ICLIST-1 | MSKELIEEIEREMDEAIEILEEMIEARRRGASEEELARLEEELKEEVRELMEKMEEAEKENNGELPKEAWPIIEEKFERFRELHREVL |
| ICLIST-2 | MSMEEKYQKWVDELVEKVMEELEKDPEYREMEDPDKWDEEKVMEIIRKVVEEERPKFEAELAA |
| ICLIST-3 | MSMEELYKKWVDKLVEKVMERLEEDPEYREMEDPDKWDEKKVMEIIREVVEEERPKFEAELAA |

**Table S2 Kinetic parameters of tandem ICLIST-2**

|  | tandem ICLIST-2 |
| --- | --- |
| *k*_a1_ (M^-1^s^-1^) | 4.78×10^5^ |
| *k*_d1_ (s^-1^) | 3.50×10^-3^ |
| *K*_D1_ (nM) | 7.32 |

Kinetic parameters were determined by fitting the sensorgrams to a bivalent analyte model. The χ^2^ value for the fit was 0.0502 RU.

**Table S3 Kinetic parameters of mutants of ICLIST-2**

|  | Q8A | R54A |
| --- | --- | --- |
| ka (M^-1^s^-1^) | 4.91×10^5^ | 2.63×10^5^ |
| kd (s^-1^) | 9.17×10^-3^ | 4.29×10^-3^ |
| *K*_D_ (nM) | 18.7 | 16.3 |

Kinetic parameters were determined by fitting the sensorgrams to a 1:1 binding model. The χ^2^ values for the fit were 0.036 and 1.47 RU.

**Table S4 Scores and amino acid sequences for ICLIST-4 to 11**

| ID | iPTM | confidence_score | Sequence | Mutated residues |
| --- | --- | --- | --- | --- |
| ICLIST-4 | 0.9627 | 0.9406 | MSLEEKYEKWVDELVEKVMEELEKDPEWGETEDPEEWDEEKVMEIIKKVVEEEKPKFEAEIAA | 10 |
| ICLIST-5 | 0.9608 | 0.9417 | MSENEKYIKWVDELIEKVMEELEKDPEFRNLEDPSKWDKEKVMEIIKKVVEEEKPKFEAELAA | 11 |
| ICLIST-6 | 0.9558 | 0.9407 | MSLEEKYEKWVDELVEKVMEELEKDPEYKNLEDPSKWDKEKVMEIIKKVVEEEKPKFEAELAA | 9 |
| ICLIST-7 | 0.9597 | 0.9409 | MSLEEKYEKWVDELVEKVMEELEKDPEYKELEDPELWDEEKVMEIIKKVVEEEKPKFEAELAA | 8 |
| ICLIST-8 | 0.961 | 0.9446 | MSDEEKYEKWVDELIEKVMEELEKDPEYKELEDPELWDEEKVMEIIKKVVEEEWPKFEAELAA | 9 |
| ICLIST-9 | 0.9582 | 0.9433 | MSLEEKYEKWVDELVEKVMEELEKDPVWGENEDPEEWDEEKVMEIIKKVVEEEKPKFEAEIAA | 11 |
| ICLIST-10 | 0.96 | 0.9422 | MSVLEKYEKWVDELIEKVMEELEKDPEWKELEDPEKWDKEKVMEIVKKVVEEEWPKFEAELAA | 12 |
| ICLIST-11 | 0.9574 | 0.9411 | MSVEEKYEKWVDELVEKVMEELEKDPEYKFLEDPEKWDKEKVMEIIKKVVEEEWPKFEAELAA | 9 |

‘Mutated residues’ indicates the number of mutations introduced into ICLIST-2

**Table S5 Hydrophobic interaction between PACAP-38 and ICLISTs**

| ICLISTs Residue | Mutation? | PACAP-38 atom | ICLISTs atom | Distance (Å) in ICLIST-2 | Distance (Å) in ICLIST-11 |
| --- | --- | --- | --- | --- | --- |
| M3 / V3 | y | 800 | 20 | N/A | 3.75 |
| Y7 | n | 765 | 58 | 3.79 | N/A |
|  |  | 797 | 58 | N/A | 3.9 |
|  |  | 798 | 56 | N/A | 3.56 |
| W10 | n | 766 | 92 | 3.23 | N/A |
|  |  | 767 | 91 | N/A | 3.33 |
| V11 | n | 740 | 99 | 3.95 | N/A |
|  |  | 765 | 100 | 3.93 | N/A |
|  |  | 742 | 98 | N/A | 3.47 |
|  |  | 766 | 99 | N/A | 3.53 |
| L14 | n | 740 | 122 | 3.99 | N/A |
|  |  | 742 | 121 | N/A | 3.57 |
| V15 | n | 680 | 131 | 3.8 | N/A |
|  |  | 709 | 132 | 3.66 | N/A |
|  |  | 740 | 132 | 3.7 | N/A |
|  |  | 742 | 131 | N/A | 3.96 |
| V18 | n | 703 | 156 | 3.89 | N/A |
|  |  | 702 | 155 | 3.83 | N/A |
|  |  | 703 | 154 | N/A | 3.63 |
| E32 | n | 590 | 279 | 3.02 | N/A |
|  |  | 673 | 277 | N/A | 3.82 |
| P34 | n | 696 | 295 | 3.58 | N/A |
|  |  | 697 | 294 | N/A | 3.41 |
| W37 | n | 671 | 326 | 3.51 | N/A |
|  |  | 670 | 328 | 3.54 | N/A |
|  |  | 696 | 327 | 3.75 | N/A |
|  |  | 703 | 328 | 3.96 | N/A |
|  |  | 731 | 325 | 3.96 | N/A |
|  |  | 672 | 326 | N/A | 3.67 |
|  |  | 671 | 328 | N/A | 3.7 |
| V42 | n | 729 | 369 | 3.47 | N/A |
|  |  | 730 | 369 | N/A | 3.72 |
| I46 | n | 741 | 402 | 3.08 | N/A |
|  |  | 726 | 403 | 3.08 | 3.31 |
|  |  | 758 | 402 | 3.79 | N/A |
|  |  | 727 | 403 | N/A | 3.09 |
|  |  | 759 | 402 | N/A | 3.8 |
| R47 | n | 757 | 409 | 3.9 | N/A |
| V50 | n | 741 | 437 | 3.74 | N/A |
|  |  | 766 | 436 | 3.52 | N/A |
|  |  | 741 | 435 | N/A | 3.37 |
|  |  | 767 | 434 | N/A | 3.36 |
| R54 / W54 | y | 767 | 475 | N/A | 3.95 |
|  |  | 798 | 476 | N/A | 3.84 |

Residues that differ between ICLIST-2 and ICLIST-11 are highlighted in yellow. ‘N/A’ indicates that no interaction was observed.

**Table S6 Salt bridge between PACAP-38 and ICLISTs**

| ICLISTs Residue | Mutation? | PACAP-38 | Distance (Å) in ICLIST-2 | Distance (Å) in ICLIST-11 |
| --- | --- | --- | --- | --- |
| Q8 / E8 | y | K20 | N/A | 4.53 |
| E23 | n | R12 / K15 | 3.64 | 3.68 |
| E32 | n | K15 | 3.25 | 3.25 |
| D33 | n | R14 | 4.57 | 4.83 |
| E35 | n | R14 | N/A | 5.3 |
| E51 | n | R30 | N/A | 5.16 |
| E58 | n | R30 | 3.6 | N/A |

Residues that differ between ICLIST-2 and ICLIST-11 are highlighted in yellow. ‘N/A’ indicates that no interaction was observed.

**Table S7 Hydrogen bond between PACAP-38 and ICLISTs**

| ICLISTs Residue | Mutation? | PACAP-38 atom | Distance (Å) in ICLIST-2 | Distance (Å) in ICLIST-11 |
| --- | --- | --- | --- | --- |
| R29 | n | 249 [O2] | 2.59 | N/A |
| E32 | n | 281 [O3] | 2.43 | N/A |
| E32 | n | 631 [O3] | 2.84 | 3.42 |

Residues that differ between ICLIST-2 and ICLIST-11 are highlighted in yellow. ‘N/A’ indicates that no interaction was observed.

**Table S8 π-stacking between PACAP-38 and ICLISTs**

| ICLISTs Residue | Mutation? | PACAP-38 atom | Distance (Å) in ICLIST-2 | Distance (Å) in ICLIST-11 |
| --- | --- | --- | --- | --- |
| Y7 | n | 796, 797, 798, 799, 800, 801 | 5.26 | N/A |
